# Modelling of SHMT1 riboregulation predicts dynamic changes of serine and glycine levels across cellular compartments

**DOI:** 10.1101/2020.12.24.424304

**Authors:** Michele Monti, Giulia Guiducci, Alessio Paone, Serena Rinaldo, Giorgio Giardina, Francesca Romana Liberati, Francesca Cutruzzolà, Gian Gaetano Tartaglia

## Abstract

Human serine hydroxymethyltransferase (SHMT) regulates the serine-glycine one carbon metabolism and plays a role in cancer metabolic reprogramming. Two SHMT isozymes are acting in the cell: SHMT1 encoding the cytoplasmic isozyme, and SHMT2 encoding the mitochondrial one. Here we present a molecular model built on experimental data reporting the interaction between SHMT1 protein and SHMT2 mRNA, recently discovered in lung cancer cells. Using a stochastic dynamic model, we show that RNA moieties dynamically regulate serine and glycine concentration, shaping the system behaviour. For the first time we observe an active functional role of the RNA in the regulation of the serine-glycine metabolism and availability, which unravels a complex layer of regulation that cancer cells exploit to fine tune amino acids availability according to their metabolic needs. The quantitative model, complemented by an experimental validation in the lung adenocarcinoma cell line H1299, exploits RNA molecules as metabolic switches of the SHMT1 activity. Our results pave the way for the development of RNA-based molecules able to unbalance serine metabolism in cancer cells.

## Introduction

The last decade has seen the rise of an RNA-centric view of many biological processes, which now flanks the more classical protein-centric view [1] [2]. The transcriptome is no longer seen just in between genome and proteome and can be regarded as a key player in many cellular processes such as differentiation, proliferation, apoptosis and many others [3] [4] [5] [6].

Many non-canonical RNA binding proteins (NC-RBPs) have been shown to interact with transcripts through novel RNA binding domains (RBDs) [3] [7]. More than 100 metabolic enzymes have been recognized as unconventional RBPs [3] [8] [9], and so far many efforts have been made to understand the hidden connections among intermediary metabolism, RNA biology and gene expression [10] [11] [12]. The non-canonical activity of numerous metabolic enzymes, also called ‘moonlighting’ [13], together with the emerging key roles of RNA molecules in several biological processes, represents a complex layer of regulation in which metabolites, RNA and metabolic enzymes co-ordinately control important biochemical pathways such as glycolysis, nucleotide biosynthesis, amino acid metabolism and many others.

Human serine hydroxymethyltransferase (SHMT) is a central enzyme of the serine-glycine-one-carbon (SGOC) metabolism. It catalyses the reversible conversion of L-serine (herein referred to as serine) into glycine, transferring a one-carbon (1C) unit to tetrahydrofolate (THF) to generate 5,10-methylentetrahydrofolate (Me-THF). Two different SHMT isoforms have been characterized in humans: SHMT1, cytosolic, and SHMT2, mitochondrial [14][15][16]. Indeed, unlike prokaryotic organisms, the SGOC metabolism is mainly compartmentalized between cytoplasm and mitochondria in all eukaryotes [17], however how the pool of enzymes, folates and amino acids are coordinately regulated as well as how the two compartments communicate is still unknown. In this work we show that the recently discovered RNA-binding ability of SHMT1 [18] links and regulates the cytoplasmic and mitochondrial SGOC metabolism, thus shaping the concentration of serine and glycine in the cell. In agreement with the evolutionary development of a compartmentalized SGOC metabolism in eukaryotes, we show that the bacterial SHMT isoform does not have a strong RNA binding propensity. We herein propose that the RNA binding property of SHMT1 has evolved to finely coordinate the cytoplasmic and micothondrial SGOC metabolism. Indeed, we show a neat example of how the RNAs are not only exploited to synthetise proteins, but also play an active functional role into the biochemical networks.

SHMT isozymes are also key players in the metabolic reprogramming of human tumours and therefore considered attractive targets for cancer therapy. However, the complex dynamic structure of the SHMT active site has so far hindered the development of effective inhibitors [19][20].

SHMT1 is a NC-RBP characterised *in vitro* and studied in lung cancer models [21] [18]. SHMT1 binds to SHMT2 mRNA through a RBD that, interestingly, partially overlaps with the folates binding cleft [18]. What is striking about such discovery is that the RNA affects the directionality of the SHMT1 reaction, favoring the cytosolic synthesis of serine rather than its cleavage [18].

Although it is already known that RNA moieties are able to inhibit or boost enzymatic reactions [22] [23], effects on the reaction equilibria have never been reported before. Exploiting a stochastic dynamic model which reliably reproduces a SHMT-centric biochemical system, the present work aims to investigate whether RNA molecules can dynamically regulate serine and glycine concentration by modulating the SHMT1 reaction. Our results show that the RNA shapes the system’s behaviour, and allows to explore multiple states in which the amounts of serine and glycine vary qualitatively. Our model proves that the SHMT1-interacting RNAs (i.e. SHMT2 mRNA and other potential interactors) link cytosolic and mitochondrial serine metabolism, fine tuning the amount of serine that can be directed into the mitochondria.

In a broader perspective, the upregulation or down-regulation of the SHMT1-interacting RNAs can represent an instrument through which the cell fine tunes serine metabolism according to its metabolic needs. Given the pivotal role of serine in the proliferation of certain types of tumors [24] [25] [26], it is conceivable that cancer cells could exploit the SHMT1-interacting RNAs to reprogram their metabolism in order to get higher levels of available serine.

## Results

Our model aims to reliably reproduce the intricate system of biochemical reactions which gives rise to the RNA-mediated crosstalk between SHMT isozymes in human cells [27] [18], thus linking cytosolic and mitochondrial serine-glycine one carbon metabolism.

Setting the SHMT isozymes at the centre of the model, we calculated the interaction propensity with RNA molecules and developed a chemical rate equations system that implements the concentration dynamic of all chemical species involved over time. Importantly, in our model we have taken into account the compartmentalization of the enzymes (cytosol and mitochondria). The diffusion among compartments is modelled as a stochastic process and rather than following the exact position of the molecules inside the cell, we consider their localization into cytosol and/or mitochondria.

### A. Protein-RNA interactions

In order to understand the evolutionary role of the RNAs in the one carbon metabolism network of eukaryotic cells, we compared the ability of the human and bacterial SHMT isoforms to bind RNAs (Supplementary Information). In Fig. 1A using *cat*RAPID [28] we have predicted the number of possible RNA binding partners of the human cytosolic and bacterial SHMT. The plot shows two types of interactions, one relative to the calculations using the bacterial RNAs and one relative to human RNAs. The human isoform clearly shows a much higher binding affinity for RNA, as confirmed *in vitro* by the electrophoretic mobility shift assay (Fig. 1B). Such observation can be interpreted as an evolutionary drift of the human enzyme towards the possibility to be regulated allosterically by RNAs (Supplementary Information).

**FIG. 1:**
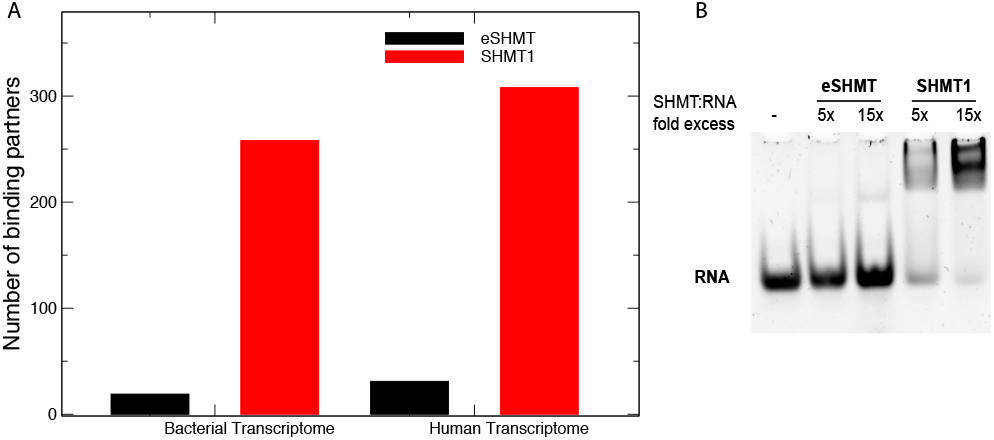
RNA binding ability of human cytosolic and bacterial SHMT. A) Bar plot of the predicted RNA interactome by *cat*RAPID relative to the human and bacterial SHMT isoforms. The predicted number of interacting RNAs of the human and bacterial SHMT are represented in red and black, respectively, the two pairs of bars are related to the calculation between the SHMTs and the human transcriptome (right) or the bacterial transcriptome (left). B) Electrophoretic mobility shift assay carried out by incubating 2 µM 50-mer RNA with 5 or 15 molar excess of bacterial SHMT (eSHMT) or human cytosolic SHMT (SHMT1).

To describe the metabolic effects caused by the recently reported RNA-mediated regulation of the SHMT1 activity [16] [18], we simulated the binding of SHMT2 mRNA to the cytosolic isozyme, taking into account the competition between RNA and folates for SHMT1 [18], Fig. 2. Using *cat*RAPID [28], we confirmed that SHMT1 binds to the 5’ untranslated region of SHMT2 (Fig. 2A), herein referred as UTR2, through a non-canonical RBD that overlaps with the folates binding cleft (N-term of the protein; Fig. 2A,B,C). Our results indicate that RNA and folate binding events are in competition. Relying on *cat*RAPID calculations [29], we could estimate the physical constants involved in the protein-RNA binding reaction (Fig. 2C), which are input of our dynamical model (Supplementary Information).

**FIG. 2:**
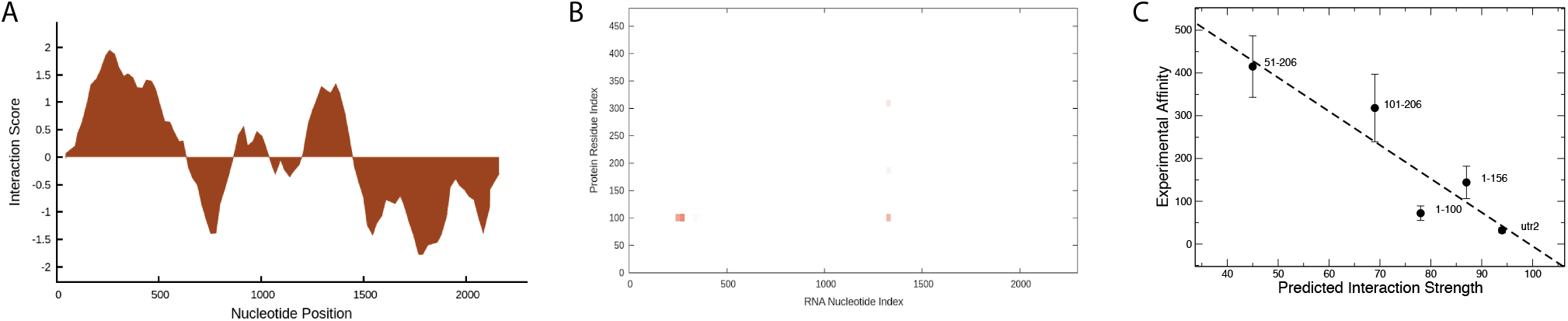
A) *cat*RAPID prediction of SHMT1 and SHMT2 mRNA interaction indicates strong binding at the 5’ UTR (UTR2). B) *cat*RAPID prediction of protein-RNA contacts indicates that the N-terminus of SHMT1, also involved in folate binding, interacts with SHMT2 mRNA in the UTR2 region. The matrix shows interaction predictions for SHMT2 nucleotides (x-axis) and SHMT1 aminoacids (y-axis). C) Prediction of SHMT1 interaction with different UTR2 fragments indicate a large variability of experimental affinities, as measured in Guiducci et al. [18].

We also have considered the SHMT1-mediated post-transcriptional regulation of SHMT2 translation. Indeed, as previously reported in [18] and [30], the cytosolic enzyme is able to downregulate the SHMT2 protein expression when overexpressed ((Supplemental figure 1)).

Worthy of note, SHMT2 mRNA plays a double role in the system: the coding region is translated into the mitochondrial enzyme, while the 5’ UTR binds to SHMT1 and modulates its activity. The double function of the SHMT2 mRNA leads the system to dynamically change the number of amino acid molecules as a function of the SHMT2 mRNA levels (which positively correlate with the mitochondrial enzyme levels).

### B. Modelling Reactions

In our model, described in Fig. 3A, the dynamic interconversion of serine and glycine, as well as of THF and Me-THF, is guided only by SHMT1 and SHMT2, according to biochemical parameters experimentally measured [16]. In the basic condition, the cytosolic enzyme catalyses the serine cleavage as well as the serine synthesis reaction, whereas the mitochondrial counterpart can only drive serine cleavage. Such asymmetry has been reported in cellular models [15] and, together with the exchange of metabolites between cytosol and mitochondria, confers a complex dynamic to the system. Cytosolic SHMT2 (i.e. SHMT2*α*) was excluded from our model as the expression is significantly lower compared to the mitochondrial isoform and its catalytic contribution can be considered negligible [18].

**FIG. 3:**
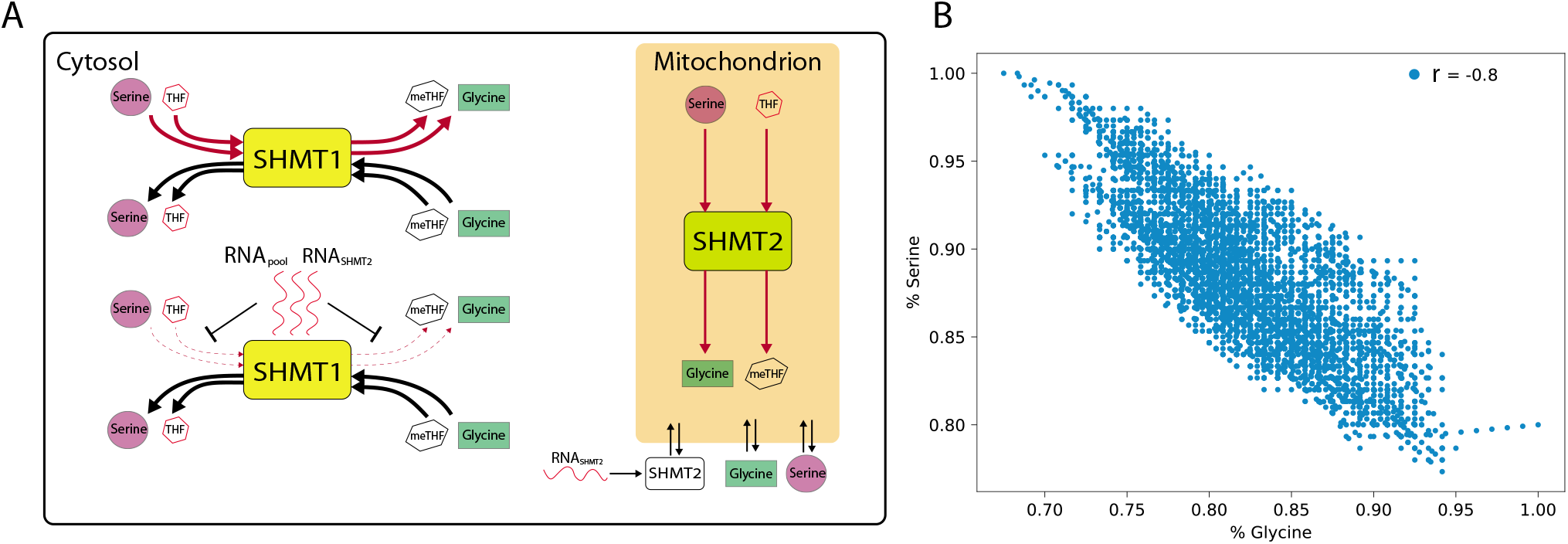
A) Scheme representing the reactions modelled in our system. The red and black arrows highlight the most favorable reactions driven by the SHMT enzymes. The effect of RNA binding on SHMT1 is to reduce the serine to glycine conversion in the cytosol (dashed red line). B) Scatter plot of serine and glycine relative free amount in solution. The plot comes from the stochastic simulation of the system without RNA. Important to notice is that the stochastic oscillations around the mean values of the concentrations are always distributed with a negative correlation (Pearson’s r = −0.8). The region of the phase space that the system explores stochastically during a simulation is finite.

Given the affinity of SHMT1 for multiple RNA moieties [18], we assumed that SHMT1, in addition to interacting with SHMT2 mRNA (3A), can bind to a wide range of RNA molecules in the cell. We built a theoretical model to explore the metabolic effects of such phenomenon: accordingly, multiple interactions have been modelled considering the presence of a cytosolic pool of RNA molecules that compete with each other for binding to the RNA binding domain of SHMT1, as seen for SHMT2 mRNA, affecting the directionality of the reaction.

The system dynamics as well as most physical constants used in our model are governed by the chemical equations derived from [16] and [18]. We have integrated the RNA molecules in the system and introduced the following dynamic to describe the effect of this new component (Supplementary Information):

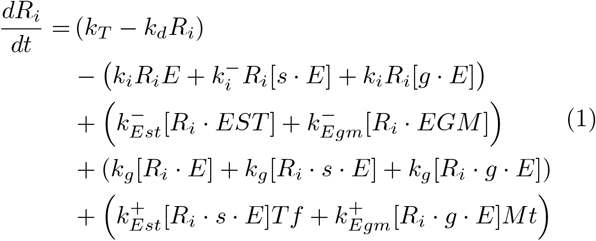

Where we have identified the RNA molecules with *R*_*i*_, SHMT1: *E*, serine: *s*, glycine: *g*, tetrahydrofolate: *Tf*, methylenetetrahydrofolate: *Mt*, the catalytic complexes SHMT1 - Serine - Tetrahydrofolate: *EST* and SHMT1 - Glycine - Methylenetetrahydrofolate: *EGM*. When represented within squared brackets, the molecules are in a complex. The first line describes the synthesis and the degradation of the RNA, we consider simply a birth-death process with no other components involved. The second line contains the terms representing the dissociation of the RNA molecule from the SHMT-substrates complexes. The third line contains terms describing the formation of the complexes with RNA molecules (Supplementary Information).

Given the spectrum of binding affinities (Fig. 2A), we considered the propensity of an RNA to interact with SHMT1 to follow a gaussian distribution centered on the average *k*_0_ associated with the experimental value measured in [18], moreover it matches very well also the predicted value from the *cat*RAPID algorithm (Supplementary Information). In this way the i-th RNA in the cytosol will bind to SHMT1 with an affinity *k*_*i*_ taken from the following Gaussian distribution:

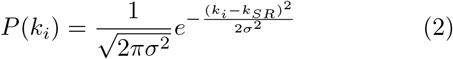

The standard deviation of the distribution *σ* has been inferred from the distribution of affinity values in Fig. 2C, however in this model the value of *σ* does not affect the results (data not shown). The biochemical reactions set in the system are described in the Supplementary Information, with a detailed table of values. All the parameters are derived from previous experiments [16] where available, whereas the missing ones have been chosen to fit experimental conditions.

We also took into account the translation of the SHMT2 mRNA; such process occurs in the cytosol and is followed by the translocation of the enzyme into the mitochondrion. We consider the transport of the molecules from and to the other cell compartment as a simple stochastic process, indeed, discerning between active and passive transport systems and looking at their effects on our biochemical model is out of the aim of this paper.

The stochastic reactions involved in the translation of the SHMT2 enzyme are the following:

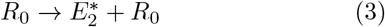

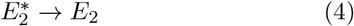

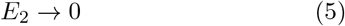

The first reaction describes the translation of the mRNA (*R*_0_). Importantly, we assume the ribosomes at a constant concentration, in order not to include their individual contribution into the model. It is important to notice that the *R*_0_ is not degraded right after its translation, implying that the concentration of *R*_0_ is not affected by the translation. The 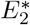 refers to the SHMT2 enzyme when still in the cytosol, then it is transported into the mitochondrion (second reaction) becoming the effective mitochondrial isoform, here called *E*_2_. The SHMT2 molecules are degraded with a rate constant. The equations that regulate the mean behaviour of the whole system, as already mentioned above, are reported in the Supplementary Information.

As schematically described in Fig.3A, the peculiarity of this biochemical network is that it evolves in parallel between different cellular compartments. Indeed, SHMT1 and SHMT2 are present in the cytosol and in the mitochondrion, respectively. As reported in [15] and schematically illustrated in Fig. 3A, SHMT1 can catalyse both serine synthesis and cleavage, whereas the mitochondrial counterpart can drive mainly the serine cleavage reaction. This asymmetry, together with the diffusion of amino acids and metabolites between cytosol and mitochondria, gives to the system a complex behaviour. As already introduced above, we quantify here the biochemical effects of the RNAs on the dynamics of the system. We aim to study the effects of the SHMT2 mRNA and the pool of RNAs present in the cytosol, understanding how such RNA species can drive the system to fine tune the amount of amino acids available in the cell.

We simulated the system using the stochastic Gillespie algorithm [31] (Supplementary Information). This well-established simulation protocol has the power to be an exact method for simulating networks of stochastic processes and it fits well the configuration of this system. This algorithm considers that all the chemical reactions depend on the concentration of the components in order to react.

Previous *in vitro* experiments have revealed the biochemical features and the dynamic of the interaction between SHMT isozymes and their substrates (either metabolites or RNA) [16][18], however their regulation in a biological context is puzzling. Most of all, it is not clear why in some cell types the SHMT1 reaction is reversible, while in others it is only directed towards serine synthesis [15].

### C. Simulation Results and Experimental Validation

In the biochemical network simulated here we consider the reactions occurring inside the cell, neglecting the transport of metabolites from the environment. To draw general conclusions, we have chosen the units of the amino acids to be considered as the percentage of free serine and glycine in the system with respect to their total concentration. Indeed, we use relative measurement also because the exact volume of the cell and its inner organelles is not easily available, thus limiting the possibility to quantify exactly the value of the amino acids concentration.

To describe how the system funnels the reaction fluxes and regulates the relative amount of amino acids, we chose to set the initial amount of serine and glycine equal to the quantity generally present in most cell mediums (i.e. serine 0.4 mM, glycine 0.4 mM) [32]. Stochastic simulations show that, in absence of RNA molecules, the amount of serine and glycine fluctuates around a mean value with negative correlation (Fig. 3B). Indeed, given the flux balance of chemical reactions, serine and glycine interchange each other in a stochastic manner, chasing an equilibrium that is impossible to keep due to the intrinsic noise of the system (Supplemental figure 2). Figure 3B shows a scatter plot describing serine and glycine concentration behaviour; of note, Pearson correlation computed between the temporal profiles of serine and glycine is *r* ≃ *−*0.8, this occurs because the system is continuously interchanging serine and glycine without any further regulation. With this setting, the biochemical system can explore a very limited space of amino acids concentration. Next step was to evaluate the effect of the SHMT1-RNA_SHMT2_ complex on the dynamics of the system. It is important to point out that the behaviour of glycine and serine reported herein refers to the amount of free molecules in the system, hence there is a fraction of amino acids that at the steady state is complexed with the other species of the system (e.g. SHMT isozymes), and it is not reported in our simulations. For the latter reason, the sum of glycine and serine appears not to be constant along the simulations.

As shown in Fig. 4A, after an initial lag phase, the levels of serine and glycine vary as a function of RNA_SHMT2_ molecules. Serine displays a bi-phasic behaviour, experiencing first a rise and then a decrease before levelling off to a constant value at high RNA_SHMT2_, while glycine level undergoes a decrease and reaches a constant amount, which does not vary further upon varying SHMT2 mRNA. This result clearly indicates that the serine/glycine levels indeed dynamically change, responding to the presence of RNA_SHMT2_ molecules, in agreement with our initial hypothesis. The increase of serine and the decrease of glycine levels have been also studied as a function of the 5’UTR of the SHMT2 mRNA (i.e. UTR2; Supplemental Figure 3A), although in the latter case the behaviour of both curves is monotonic.

**FIG. 4:**
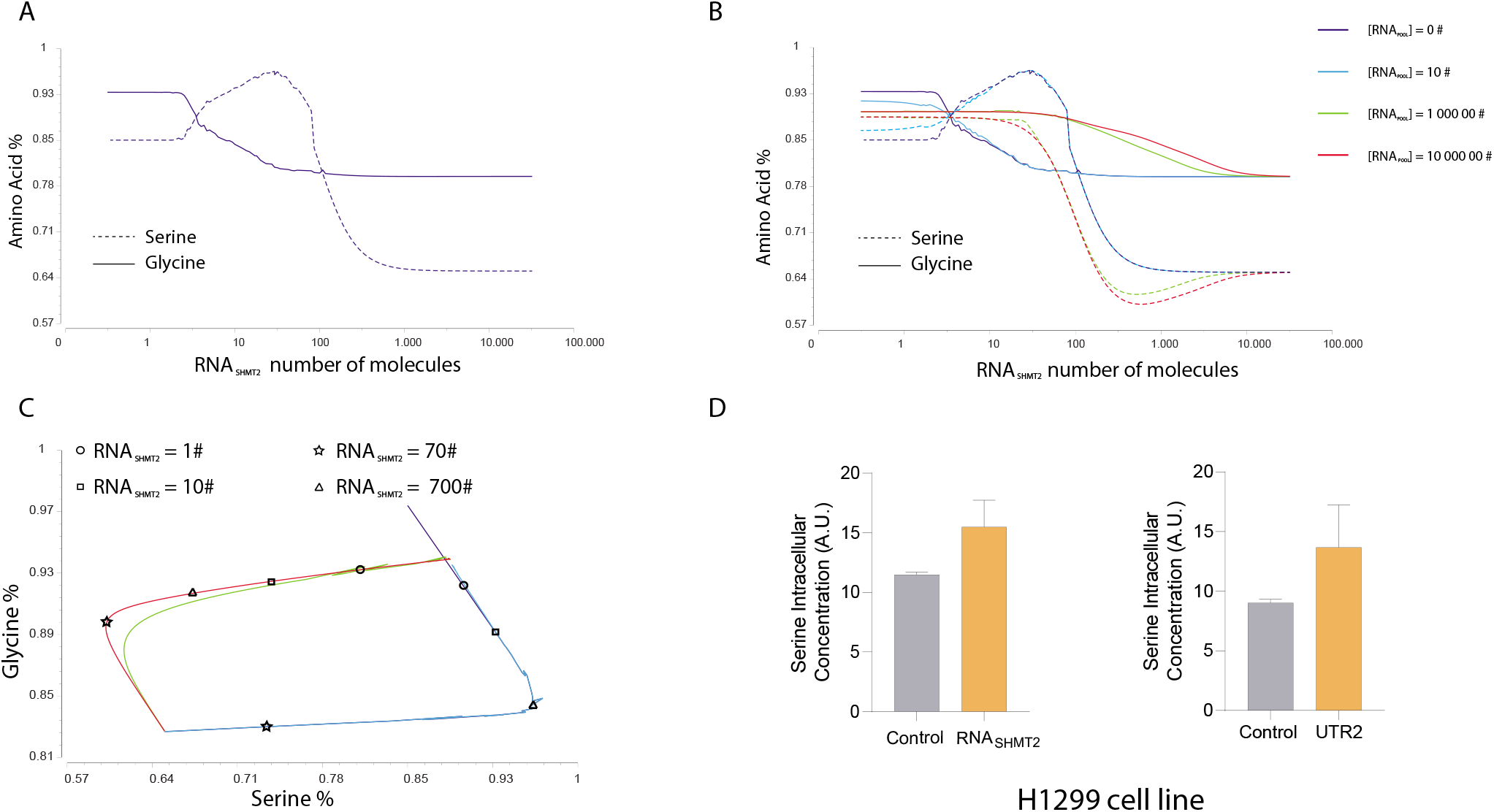
Behaviour of the final steady state of the system for different amounts of RNA molecules inserted in the simulations. A) Behaviour of serine and glycine as a function of the SHMT2 mRNA (RNA_SHMT2_), considering the RNA pool equal to 0. B) Same dynamic described in A but considering different numbers of RNA pool (RNA_pool_) molecules present in the system. C) Phase space of glycine and serine for the final steady states. The plot represents the values reported in panel B but on the serine-glycine plane. Each curve represents the amount of serine and glycine concentrations at the steady state in the system, each point is calculated for different values of RNA_SHMT2_. The amounts of RNA_SHMT2_ computed for each region of the curves are indicated with symbols. The colours of the curves indicate the different values of RNA_pool_ used in each simulation, the legend is in common with panel B. It is important to notice that the accessible region of the pair of values glycine-serine can change dramatically for different values of RNA_pool_. Indeed, by changing RNA_pool_ in the system the amino acids explore couples of values not possible to access by only modulating RNA_SHMT2_. It is possible to appreciate how different classes of RNAs can drive the system to explore different ranges of serine and glycine concentrations. D) H1299 were transiently transfected with a control empty vector (Control), the UTR2-shmt2 expression-inducing vector (RNA_SHMT2_) or the UTR2 sequence expression-inducing vector (UTR2) and, after 24 hours, treated with tetracycline 0.01 mg/ml in complete RPMI medium. 24 hours after the treatment, cells were harvested, processed and gas chromatography/mass spectrometry was used to quantify the intracellular serine content. Data derive from two independent experiments.

Considering that SHMT1 will have multiple target RNAs in the cytoplasm, we simulated the presence of other RNAs interacting with the enzyme; such assumption implies that the pool of RNAs will compete with folates substrates as well, contributing to inhibit the serine cleavage reaction while slightly affecting serine synthesis. We investigated if and how the presence of these RNA moieties (named RNA_pool_ in Fig. 4B) affects amino acid levels. Fig. 4B shows how serine and glycine amounts change as a function of RNA_SHMT2_ molecules in the system, different coloured curves are then referred to different values of RNA_pool_ molecules considered in the simulation. As before, we quantified the amount of amino acids present in the system as a percentage of their maximum availability.

We identified two regimes, relative to low (dark and light blue curves in Fig. 4B) and high (red and green curves in Fig. 4B) amount of RNA_pool_ molecules. Each point of the curves represents the mean value of the steady state amino acids concentration over different simulations. In the first couple of curves (i.e. light and dark blue), we can see again the very interesting and non-monotonic behaviour of the serine and glycine concentration, also seen in the absence of RNA_pool_ (Fig. 4A). When the amount of RNA_pool_ is high (red and green curves), a more symmetric system with similar starting levels of serine and glycine is observed. In Fig. 4C it is plotted the mean percentage of serine and glycine for a given amount of RNA_SHMT2_ and RNA_pool_. Being a different representation of the data already plotted in Fig. 4B, such curves aim to drive the attention on a different aspect of the system. Indeed, they highlight the fact that varying the RNA molecules in the system, it is possible to explore and sample a wide region of the phase space of glycine and serine concentrations. Worthy of note, moving through this space in a controlled manner implies that the cell could be able to regulate the relative amino acid concentrations simply by modulating the amount of RNAs bound to SHMT1. For instance, for low amounts of RNA_pool_ is possible to have an higher percentage of free serine combined with a lower concentration of glycine, on the other hand, for high values of RNA_pool_ molecules, the system explores a lower serine concentration combined with a higher amount of glycine.

The results of the simulations were validated in the lung adenocarcinoma cell line H1299, for which the SHMTs crosstalk has been previously described [18]. In our experiments, we exploited an inducible vector to over-express SHMT2 mRNA (i.e. RNA_SHMT2_ involving the 5’UTR and the coding region) and we measured serine and glycine intracellular concentrations upon transfection (see Supplemental Figure 4 and Methods for the description of the inducible system). An increase in the serine concentration has been observed upon overexpression of such construct with respect to the control (Fig. 4D), thus reproducing the increase simulated in Fig. 4A; on the other hand, we could not observe variations in glycine levels (Supplemental Figure 3D). The latter observation suggests that intracellular glycine homeostasis results from a more complex balance between uptake and utilization/formation by several metabolic pathways, as previously observed [33] [34]. Finally, similar results have been obtained overexpressing only the 5’UTR of the construct (i.e. UTR2), Fig. 4D and Supplemental Figure 3B,C. These results fully agree with previous data obtained on the HAP1-SHMT2KO cell line (Supplemental Figure 5), confirming the general validity of the effect of the UTR2 on SHMT1 activity [18], as predicted by the model.

## Discussion

The integration of an alpha-proteobacterial endosymbiont in the cell [35] allowed the development of a complex compartmentalized metabolic network that requires the interplay of enzymes present in mitochondria and cytoplasm to fine tune metabolites availability [36]. L-serine, glycine and folates are the key metabolites of the SGOC metabolism and are uptaken by the cell from the environment [25, 26, 37]. In addition, serine and glycine can be synthetized *de novo* through the serine synthesis pathway and from threonine [25, 38], respectively, and they can be interconverted thanks to SHMT enzymes [15, 39]. Here we used state-of-the-art theoretical approaches to demonstrate that RNA molecules link the activities of human cytosolic serine hydroxymethyl-transferase (SHMT1) and its mitochondrial counterpart (SHMT2). The dynamic modulation of serine and glycine orchestrated by SHMT enzymes is fundamental for cell growth and, together with the enhancement of pathways generating such aminoacids [26, 40], plays an important role in cancer [25]. Intriguingly, while bacterial SHMT does not have RNA binding ability (Fig. 1), human cytosolic SHMT1 has evolved the ability to interact with RNA [18]. The biochemical effects of RNA interactions with SHMT1 and their impact on SHMT2 activity represent one of the missing links between cytosolic and mitochondrial SGOC metabolism, having important repercussions on cell proliferation. Suffice to say that mitochondrial SHMT2 and the immediately downstream mitochondrial enzyme 5,10-methylene-tetrahydrofolate dehydrogenase (MTHFD2) are the most consistently over-expressed metabolic enzymes in cancer [41]. Thus, it is possible to speculate that the choice of using an already existing element of the interaction networks (i.e. RNA_SHMT2_), without the need to recruit another protein to perform an allosteric regulation of human SHMT1, can be seen as a sign of optimization driven by evolution.

Indeed, the data presented herein suggest that an RNA-mediate control of amino acid concentrations is operative inside the cell, exerted through the moonlighting activity of human cytosolic SHMT1. The cell can explore a different range of amino acids concentration by tuning the expression of RNAs interacting with the cytosolic enzyme SHMT1, acting as a sensor of the metabolic status of the cell.

On the one side, RNA_SHMT2_ is able to allosterically modulate SHMT1 activity, by binding through its 5’UTR and thus selectively inhibiting serine cleavage. On the other side, RNA_SHMT2_ is translated into the mitochondrial enzyme which controls serine to glycine conversion in the organelle. The combined action of the sensor (SHMT1) and the multifunctional RNA effector (RNA_SHMT2_) allows the system to access states of serine and glycine concentrations impossible to explore otherwise.

For low SHMT2 mRNA levels, serine and glycine amounts are similar, with a slightly higher amount of glycine due to the fact that the system has an intrinsic asymmetry (Fig. 4A). As soon as SHMT2 mRNA levels increase, serine raises to reach a maximum, whereas glycine levels lower. Remarkably, this dynamic accurately describes the modulation of serine levels experimentally determined in the leukaemia HAP1 cells containing a frameshift mutation in the RNA_SHMT2_ and thus lacking the synthesis of the SHMT2 protein [18] (Supplemental Figure 5), and it has been further confirmed here in the lung adenocarcinoma cell line H1299. Indeed, by increasing SHMT2 mRNA levels, an increase in serine intracellular levels has been measured, in agreement with predictions (Fig. 4D). The overexpression of the 5’ UTR alone, which corresponds to the SHMT2 mRNA region exerting the allosteric effect, increases the levels of intracellular serine (Fig. 4D); under these conditions, we previously demonstrated that SHMT1 serine cleavage is indeed inhibited [18] and thus intracellular serine is consumed less, well in agreement with the results of the simulation. In a wild type background, where the translation of RNA_SHMT2_ in the functional mitochondrial protein acts coordinately with the inhibitory effect of the same mRNA, the accumulation of serine (Fig. 4A) is the result of an insufficient amount of mitochondrial enzyme that transforms the substrate into glycine. Thus, in the tested range of RNA values, the main metabolic effect is the inhibition of the SHMT1 mediated serine cleavage. As the amount of RNA_SHMT2_ raises, the RNA binding site of SHMT1 becomes saturated, and more SHMT2 mRNA molecules become available to be translated into the mitochondrial protein, which is catalytically active in the mitochondria and thus starts to affect serine levels; indeed, as expected, serine starts to be rapidly consumed, as shown by the drop right after the peak (Fig. 4A). Importantly, glycine does not decrease further, because the saturation effect driven by the RNA mediated inhibition of SHMT1 is compensated by a higher concentration of mitochondrial enzyme, which converts serine into glycine. Finally, when the levels of SHMT2 mRNA are extremely high, the translation of mitochondrial enzyme reaches a plateau, explaining why the serine curve gets stabilized.

In the presence of a more complex population of allosteric RNA effectors of SHMT1 (Fig. 4B), the intermediate serine peak disappears because SHMT1 is synergistically saturated by RNA_SHMT2_ and RNA_pool_. Interestingly, under these conditions, the amino acids concentration remains constant for a wide range of RNA_SHMT2_, until the point in which both serine and glycine start to drop. The increase of RNA_SHMT2_ causes the increase of SHMT2 in the mitochondria, which indicates that the SHMT2 reaction becomes favoured. The pronounced drop of serine can be explained by the fact that the reaction driven by SHMT2 consumes serine more quickly as the amount of enzyme increases; in addition, a significative amount of serine is bound to the cytosolic RNA-protein complexes and thus is not detected by our model. In addition, in the presence of a pool of RNAs, these molecules behave as competitors of RNA_SHMT2_ for the binding to SHMT1; a left shift of the curve is observed in the region where serine levels drop with respect to the behaviour recorded in the absence of the RNA_pool_ (Fig. 4B). Such phenomenon can readily be explained by envis-aging that the number of free RNA_SHMT2_ molecules able to yield active SHMT2 enzyme is higher under these conditions, allowing serine to be more efficiently consumed. In the final part of the curve, a critical concentration of RNA_SHMT2_ is reached, where the expression of the mitochondrial protein is saturated by the finite number of ribosomes set in the system. Indeed, the mitochondrial reactions are saturated as well, whereas in the cytosol the increasing levels of RNA_SHMT2_ will increase inevitably the amount of amino acids bound into complexes. Thus, glycine starts to compete with serine complexed with SHMT1 and RNA, letting the amount of free serine to increase until it reaches the equilibrium.

In summary, tuning amino acids concentrations is crucial for the cell behaviour and fate, and we showed that RNA can act as key regulator in this scenario. These predictions and observations offer an unprecedented and perspective view of how RNA dynamics might directly control metabolite dynamics through key protein sensors. RNA binding proteins are well known to regulate RNA metabolism, function and translation; a novel mechanism has been recently described for the RNA-mediated regulation of proteins function, referred to as riboregulation [42]. Intriguingly, many of the proteins classified as non-conventional RNA binding proteins not only are indeed metabolic enzymes [8, 43–45], but also have isoforms localized in different subcellular compartments, such as SHMT in the present paper, which might undergo a differential allosteric regulation by RNA molecules depending on their location. The advancement of aptamer-based therapeutics in oncology [46, 47], combined with the recent evidences of riboregulation of metabolic enzymes involved in cancer [3, 10, 11, 18, 45], open the possibily to employ RNA aptamers to regulate metabolic enzymes and lower their oncogenic potential. The selectivity of such RNA modulators can be tailored to hit specific cell types, taking advantage of the ongoing knowledge on cell-specific targeting of RNA molecules employing different mechanisms of action [48], [49]. In this view, our results pave the way for the design of new aptamers-based treatments employing RNA molecules as allosteric modulators of the activity of selected metabolic enzymes, such approach will be helpful to treat diseases where the dysregulation of serine and glycine plays an important role, including cancer.

## Material and methods

We have used the COPASI software [31] in order to implement the Gillespie algorithm for the stochastic simulations. Protein-RNA binding predictions have been computed using the *cat*RAPID software [28]. The information related to the electrophoretic mobility shift assay and serine/glycine intracellular measurements are provided in the Supplementary Informations.

## Acknowledgments

We acknowledge Alexandros Armaos for fruitful discussion and for the *cat*RAPID guidelines and Alberto Macone for the assistance with the GC/MS experiments. The research leading to these results has been supported by European Research Council (RIBOMYLOME 309545 to GGT and ASTRA 855923 to GGT), the H2020 projects IASIS 727658 and INFORE 25080, the Spanish Ministry of Economy and Competitiveness BFU2017-86970-P as well as the collaboration with Peter St. George-Hyslop financed by the Wellcome Trust. GGu is supported by the Barts Charity Grant. Funding from AIRC under IG2019-ID23125 project– PI FC and from Sapienza University of Rome (Grants n. RG11816430AF48E1, RM11916B46D48441, RP11715C644A5CCE, GA116154C8A94E3D to F.C.) are gratefully acknowledged.

## Author contributions

MM and GGu conceptualized the theoretical approach. Validation: AP, SR, FRL and GGi designed and performed the experiments. GGT implemented the catRAPID analysis. Supervision: MM developed the metabolic model with the help of GGu supervised by GGT. FC designed and supervised the experiments. Writing: MM, GGu, GGT and FC wrote the paper. Visualization: MM, GGT, GGu, GGi took care of data. Funding acquisition: GGT and FC. MM and GGu are to be considered co-first authors. To whom correspondence may be addressed. E-mail: MM and GGT for the computational parts; FC for the experimental part.

