## Supplementary Materials for "Modelling of SHMT1 riboregulation predicts dynamic changes of serine and glycine levels across cellular compartments"

### Supporting Information

This document contains the supporting info for the paper "Modelling of SHMT1 riboregulation predicts dynamic changes of serine and glycine levels across cellular compartments". Here we provide supplemental figures and experimental procedures, as well as the tables containing the chemical reactions used in the model with the relative constants. Moreover, to give a complete view of the model, we write down the mathematical differential equations that regulate the mean behavior of the system.

#### I. DATA GENERATION, THE GILLESPIE ALGORITHM

The Gillespie algorithm is an exact stochastic simulation procedure for a set of Poissonian events connected among each other. The algorithm, introduced by D.J. Gillespie in 1970 [1], is very good to simulate well mixed systems of chemical reactions. Indeed, in the computation the diffusion is neglected considering that at each time step all the molecules can be in contact with each other. In this context it is considered that each reaction of the system follows the Arrenius law, so the probability of a reaction $j$ happen in a time $\tau$ will be:

$$P(j,\tau) = e^{-a_j \tau} \tag{1}$$

where $a_j$ is the propensity of the reaction $j$, that in the linear case will simply be $k_j x_j$ where the $k_j$ and $x_j$ are the rate and substrate of the reaction rate. Considering a set of independent processes the probability of having a certain reaction after time $\tau$ will be the product:

$$P(\tau) = \prod_j e^{-a_j \tau} = e^{-\sum_j a_j \tau} = e^{-a_{tot} \tau} \tag{2}$$

where is introduced $a_{tot} = \sum_j a_j$ the total propensity to react of the system. This defines an exponential probability distribution of having one reaction after time $\tau$. The next step is to define which is the reaction that will fire, this is taken simply from the relative propensity of each reaction:

$$P(j) = \frac{a_j}{a_{tot}} \tag{3}$$

Summarizing it is possible to recapitulate the steps of the algorithm as the follow:

1. Define the Network, Matrix of interaction, nature of interactions, rate of the reactions, number of species (nodes), final time $T_{max}$ of the simulation and propensity functions $a_j$.
2. Select the time of the first reaction from the exponential distribution of probability of the times: $P(\tau) = e^{-a_{tot}\tau}$
3. Select the reaction from the relative propensity: $P(j) = \frac{a_j}{a_{tot}}$
4. upgrade the species according to the rules of the reaction $j$
5. compute the new propensities $a_j$ and $a_{tot} = \sum_j a_j$ from the updated set of the network.
6. update the time $t += \tau$
7. go to step 1 and iterate until $t < T_{max}$

For a system well mixed system where the statistical structure does not take into account diffusion but just the reaction rates this algorithm gives exactly the real time traces of the status of the chemical species involved. It is important to notice that the time step of the simulation is not constant, but being simulation of a Markov process, where the state of the net at time $T + \tau$ depends just on the state a time $t$ we can force the print out the state of the network for any $dt$.

#### II. THE CATRAPID ALGORITHM

The *cat*RAPID algorithm [2] estimates the binding potential through van der Waals, hydrogen bonding and secondary structure propensities of both protein and RNA sequences allowing identification of binding partners with high confidence. Indeed, as reported in a recent analysis of about half a million of experimentally validated interactions [3] the algorithm is able to separate interacting vs non-interacting pairs with an area under the ROC curve of 0.78 (with False Discovery Rate FDR significantly below 0.25 when the Z-score values are $> 2$) .

*cat*RAPID omics [4] was used to compare RNA interactions of human and bacterial SHMT. Different sets of protein-RNA interactions were calculated using the overall score (cut-off of 0.5).

Human SHMT1 vs human transcriptome:
`http://crg-webservice.s3.amazonaws.com/submissions/2020-11/310289/output/all_interactions.310289.%3Esp%7CP34896%7CGLYC_HUMAN_.txt`

Bacterial SHMT vs human transcriptome:
`http://crg-webservice.s3.amazonaws.com/submissions/2020-11/310288/output/all_interactions.310288.sp%7CP0A825%7CGLYA_ECOLI.txt`

Bacterial SHMT vs bacterial transcriptome (divided in 5 parts):
`http://crg-webservice.s3.amazonaws.com/submissions/2020-12/313771/output/index.html?unlock=a2bfc632b5`
`http://crg-webservice.s3.amazonaws.com/submissions/2020-11/311729/output/index.html?unlock=613b8dd744`
`http://crg-webservice.s3.amazonaws.com/submissions/2020-11/311730/output/index.html?unlock=6ea415adc7`
`http://crg-webservice.s3.amazonaws.com/submissions/2020-11/311612/output/index.html?unlock=87d0f6cb85`
`http://crg-webservice.s3.amazonaws.com/submissions/2020-11/311731/output/index.html?unlock=18b53a2565`

Human SHMT1 vs bacterial transcriptome (divided in 5 parts):
`http://crg-webservice.s3.amazonaws.com/submissions/2020-11/312124/output/index.html?unlock=e676dbf4ed`
`http://crg-webservice.s3.amazonaws.com/submissions/2020-11/312125/output/index.html?unlock=f1aee6e225`
`http://crg-webservice.s3.amazonaws.com/submissions/2020-11/312126/output/index.html?unlock=7f647ec3b4`
`http://crg-webservice.s3.amazonaws.com/submissions/2020-11/312127/output/index.html?unlock=52ec623679`
`http://crg-webservice.s3.amazonaws.com/submissions/2020-11/312128/output/index.html?unlock=a01695a0ac`

The *cat*RAPID fragment approach [5] was used to predict SHMT1−SHMT2 interactions:
`http://crg-webservice.s3.amazonaws.com/submissions/2020-11/305872/output/index.html?unlock=b39930694e`

The binding constant Kd was estimated using catRAPID strength [6]:
wt: `http://service.tartaglialab.com/email_redir/160155/d0759bf3a1`
51-206 `http://service.tartaglialab.com/email_redir/160124/5c284c856d`
1-156 `http://service.tartaglialab.com/email_redir/160123/30cfd85a7a`
1-100 `http://service.tartaglialab.com/email_redir/160121/5c3c0d26f5`
101-206 `http://service.tartaglialab.com/email_redir/160119/5abb140cb0`

#### III. SUPPLEMENTAL RESULTS AND EXPERIMENTAL PROCEDURES

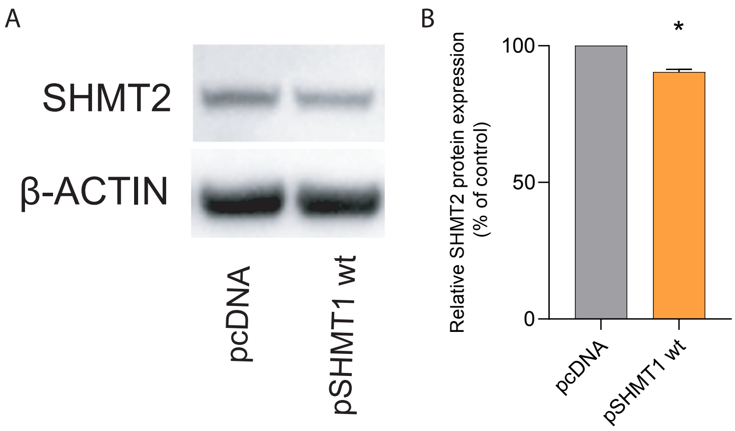


FIG. 1: Effects of SHMT1 overexpression on SHMT2 protein levels. A) Data taken from [7]. SHMT2 protein expression analysed by western blot in the A549 cell line transfected with an empty vector (pcDNA) or with a vector overexpressing SHMT1 (pSHMT1 wt). B) Densitometric analysis of the western blot showed in panel A.

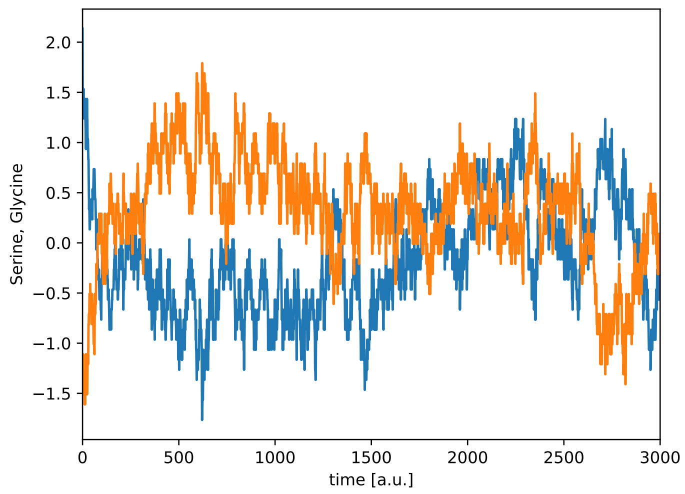


FIG. 2: Time traces of serine (blue) and glycine (orange) from the numerical simulation without RNA molecules present in the system. The values plotted are the differences between the mean value of the simulation and the current value of the concentration. We have used this sub-model in order to match the physical constants that are not available in literature. It is important to notice that the amino acids have a chasing dynamic, where each positive fluctuation of one amino acid correspond to a negative fluctuation of the other.

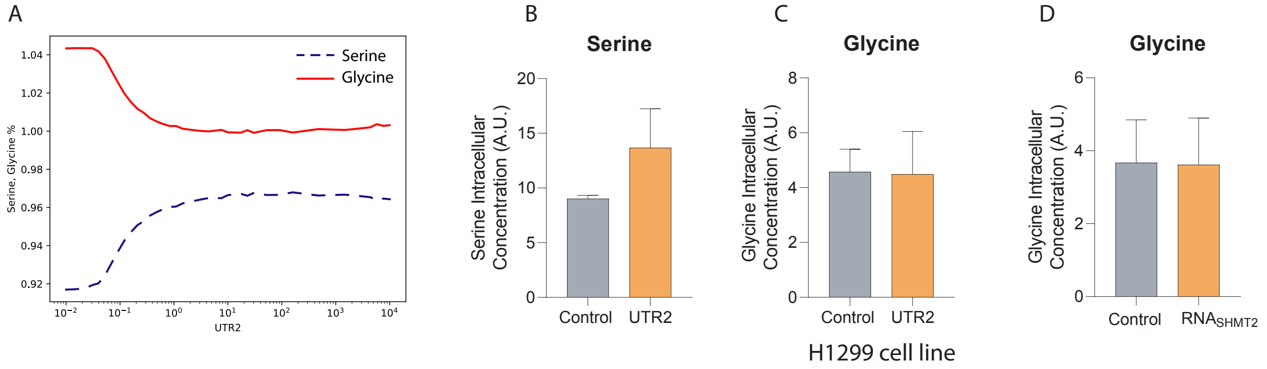


FIG. 3: A) Final steady state of serine and glycine as a function of the isolated UTR2 RNA number of molecules. The model predicts that in the absence of the SHMT2 coding sequence (and translation) the decrease in serine levels due to the activity of the mitochondrial enzyme (see Figure 4A in main text) should not be observed and, accordingly, both serine and glycine follow a monotonic behaviour. B) Serine and C) glycine intracellular concentration measured in H1299 upon transfection of an empty vector (Control) or a vector for the inducible expression of UTR2. The expression of UTR2 increases serine but not glycine levels. D) Glycine intracellular concentration measured in H1299 upon trasfection of an empty vector (Control) or a vector for the inducible expression of the SHMT2 mRNA ($RNA_{SHMT2}$). The expression of $RNA_{SHMT2}$ does not cause changes in intracellular glycine levels. All experiments have been performed in duplicate.

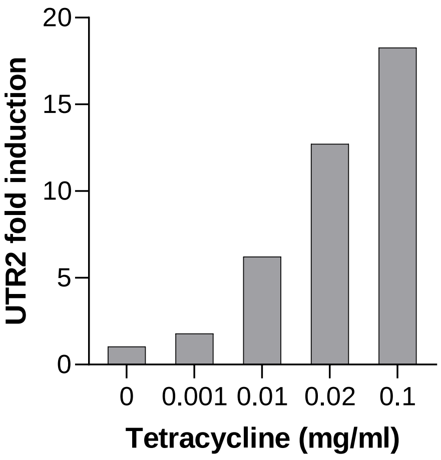


FIG. 4: UTR2 expression analysed by qRT-PCR in the H1299 cell line transfected with a vector overexpressing different levels of UTR2 as a function of tetracycline concentration. Data derive from a single replicate. The increase of tetracycline concentration induces the transcription of higher amount of the UTR2 sequence with 0.8-, 6-, 12- and 18-fold induction with 0.001, 0.01, 0.02, 0.1 mg/ml tetracycline, respectively.

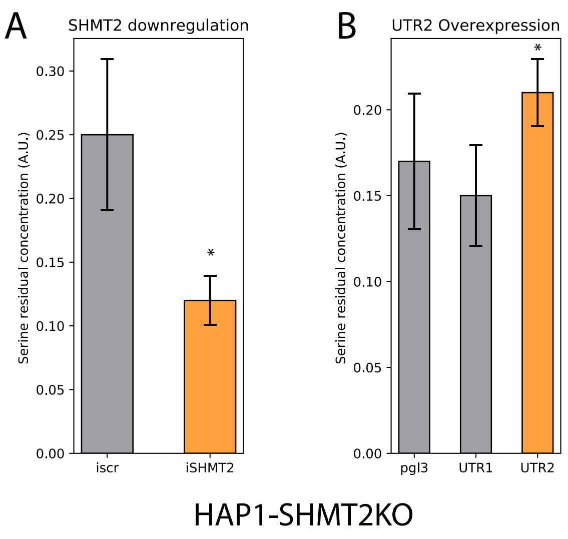


FIG. 5: Evaluation of serine residual concentration in the growth medium of the HAP1SHMT2KO line, which expresses SHMT2 mRNA but not its protein product. Data taken from [8]. A) Serine levels in the extracellular medium were measured 72 h after transfection with RNAi to knockdown the SHMT2 mRNA (iscr, scramble RNA; iSHMT2, shmt2 interference) or B) with plasmids to overexpress UTR2 (pgl3, empty vector; UTR1, UTR1-containing plasmid; UTR2, UTR2-containing plasmid). Statistical analyses are performed on three independent experiments, $*P < 0.05$.

**Electrophoretic mobility shift assays.** SHMT from Escherichia coli (eSHMT) (a kind gift from Prof. Contestabile, Sapienza University of Rome) was used as a reference for prokaryotic SHMTs; human cytosolic SHMT (SHMT1), purified as reported in [8], was used as a control. eSHMT was quantified according to [9]., 2007. RNA used for binding assay is a 50-fragment of the 206-mer used in [8]. Binding reaction was run at RT by incubating 2 µM RNA with different molar excess of protein (5x or 15x) for 20 min in 12 µl of binding buffer (20 mM HEPES pH 7.4, 100 mM NaCl) containing 20 µg/ml bovine serum albumin (BSA) and 8% (v/v) glycerol. The reaction mixtures were separated onto 4% nondenaturing polyacryleamide gels in 0.5× TBE buffer (45 mM Tris-Borate, 1 mM ethylenediaminetetraacetic acid, pH 8.6) run at 80 Volts for 30 min. For the visualization of RNA fragments, gels were stained with SYBR Safe (Invitrogen) in 30 ml of 0.5× TBE; images were acquired using Chemidoc MP Imaging System (Bio-Rad). The experiment was run in duplicate.

**Western blot analysis.** Experiments were taken from [7].

**Cell line and transfections.** H1299 cancer cells from ATCC (Manassas, VA, USA) were maintained in RPMI medium supplemented with 2 mM L-glutamine, 100 IU/ml penicillin/streptomycin, and 10% fetal calf serum (FCS; Sigma-Aldrich, St Louis, USA). H1299 cells were transfected using jetPRIME transfection reagent (Polyplus transfection, New York, USA) with the epB-Puro TT tetracycline inducible empty vector kindly provided by Dr. Fatica (Sapienza University of Rome), the same vector modified to induce the expression of the UTR2 sequence or the same vector inducing the expression of the UTR-shmt2 sequence were provided by Genscript Biotech (Leiden, Netherland). 24 hours after transfection the cells were treated with 0.01 mg/ml tetracycline (Sigma-Aldrich).

**RNA extraction and real time qRT-PCR.** 24 hours after tetracycline treatment the cells were harvested and washed twice with PBS, RNA was extracted and purified using TRIzol reagent (Invitrogen, Carlsbad, USA) following the manufacturer instruction. 1 ug of the purified RNA was retrotranscribed using SuperScript First-Strand Synthesis System Kit (Invitrogen) following the manufacturer instruction. The qRT–PCR analysis performed in triplicate for each sample using Power-Up sybr green master mix (Applied Byosystems, Austin, USA) following the manufacturer instructions. Reactions were performed using Quant Studio 3 (Applied Biosystems). Primers used were previously published in [8].

**Serine/glycine content analysis.** 24 hours after tetracyclin treatment the cells were harvested, washed twice in PBS and the samples were extracted and derivatized for serine and glycine intracellular content analysis [10] using gascromatography/mass spectrometry as previously described in [8].

**Serine and glycine consumption assay**. Experiments in HAP1 cells were taken from [8].

#### IV. ODE OF THE SYSTEM

In this section we listed all the differential equations that regulate the mean behavior of the system. An analytical solution of such a huge system clearly is not possible to achieve. In this set of differential equation is also encoded the nature of chemical reaction involved and its rate law.

$$
\begin{aligned}
\frac{\mathrm{d}([\mathrm{E}] \cdot V_{\mathrm{Cell}})}{\mathrm{d}t} = & -V_{\mathrm{Cell}} \cdot \left(\frac{[\mathrm{E}] \cdot \mathrm{k\_cat} \cdot [\mathrm{Ser}]}{[\mathrm{Ser}] + \mathrm{Km}_{(\mathrm{reaction\_0})}}\right) \\
& -V_{\mathrm{Cell}} \cdot \left(\frac{[\mathrm{E}] \cdot \mathrm{k\_cat}_{(\mathrm{reaction})} \cdot [\mathrm{THF}]}{[\mathrm{THF}] + \mathrm{Km}_{(\mathrm{reaction})}}\right) \\
& -V_{\mathrm{Cell}} \cdot \left(\frac{[\mathrm{E}] \cdot \mathrm{k\_cat} \cdot [\mathrm{Gly}]}{[\mathrm{Gly}] + \mathrm{Km}_{(\mathrm{reaction\_9})}}\right) \\
& -V_{\mathrm{Cell}} \cdot \left(\frac{[\mathrm{E}] \cdot \mathrm{k\_catC} \cdot [\mathrm{Met}]}{[\mathrm{Met}] + \mathrm{Km}_{(\mathrm{reaction\_10})}}\right) \\
& +V_{\mathrm{Cell}} \cdot \left(\mathrm{k1}_{(\mathrm{reaction\_11})} \cdot [\mathrm{MetE}]\right) \\
& +V_{\mathrm{Cell}} \cdot \left(\mathrm{k1}_{(\mathrm{reaction\_12})} \cdot [\mathrm{GlyE}]\right) \\
& -V_{\mathrm{Cell}} \cdot (\mathrm{kg} \cdot [\mathrm{RNA}] \cdot [\mathrm{E}]) \\
& +V_{\mathrm{Cell}} \cdot (\mathrm{kg} \cdot [\mathrm{RNA\_E}]) \\
& +V_{\mathrm{Cell}} \cdot \left(\mathrm{k1}_{(\mathrm{reaction\_3})} \cdot [\mathrm{SerE}]\right) \\
& +V_{\mathrm{Cell}} \cdot \left(\mathrm{k1}_{(\mathrm{reaction\_4})} \cdot [\mathrm{EThf}]\right) \\
& -V_{\mathrm{Cell}} \cdot (\mathrm{kfull} \cdot [\mathrm{fRNA}] \cdot [\mathrm{E}]) \\
& +V_{\mathrm{Cell}} \cdot (\mathrm{kfull} \cdot [\mathrm{fRNA\_E}])
\end{aligned}
$$

$$\frac{\mathrm{d}([\mathrm{Ser}]\cdot V_{\mathrm{Cell}})}{\mathrm{d}t} = -V_{\mathrm{Cell}} \cdot \left(\frac{[\mathrm{E}]\cdot \mathrm{k\_cat}\cdot [\mathrm{Ser}]}{[\mathrm{Ser}] + \mathrm{Km}_{(\mathrm{reaction\_0})}}\right)$$
$$-V_{\mathrm{Cell}} \cdot (\mathrm{kInib} \cdot [\mathrm{MetE}] \cdot [\mathrm{Ser}])$$
$$+V_{\mathrm{Cell}} \cdot (\mathrm{kInib} \cdot [\mathrm{ESM}])$$
$$-V_{\mathrm{Cell}} \cdot \left(\frac{[\mathrm{EThf}]\cdot \mathrm{k\_cat}\cdot [\mathrm{Ser}]}{[\mathrm{Ser}] + \mathrm{Km}_{(\mathrm{reaction\_2})}}\right)$$
$$-V_{\mathrm{Cell}} \cdot (\mathrm{kg} \cdot [\mathrm{RNA\_E}] \cdot [\mathrm{Ser}])$$
$$+V_{\mathrm{Cell}} \cdot \left(\mathrm{k1}_{(\mathrm{Rna\_10})} \cdot [\mathrm{RnaSerE}]\right)$$
$$+V_{\mathrm{Cell}} \cdot \left(\mathrm{k1}_{(\mathrm{reaction\_3})} \cdot [\mathrm{SerE}]\right)$$
$$-V_{\mathrm{Cell}} \cdot (\mathrm{K\_diff\_in} \cdot [\mathrm{Ser}])$$
$$+V_{\mathrm{Cell}} \cdot (\mathrm{k\_diff\_out} \cdot [\mathrm{Serm}])$$
$$+V_{\mathrm{Cell}} \cdot \left(\mathrm{k1}_{(\mathrm{reaction\_5})} \cdot [\mathrm{EST}]\right)$$
$$+V_{\mathrm{Cell}} \cdot (\mathrm{alphaS})$$
$$-V_{\mathrm{Cell}} \cdot (\mathrm{betaS} \cdot [\mathrm{Ser}])$$
$$-V_{\mathrm{Cell}} \cdot (\mathrm{kfull} \cdot [\mathrm{fRNA\_E}] \cdot [\mathrm{Ser}])$$
$$+V_{\mathrm{Cell}} \cdot (\mathrm{kfull} \cdot [\mathrm{fRnaSerE}])$$

$$\frac{\mathrm{d}([\mathrm{Met}]\cdot V_{\mathrm{Cell}})}{\mathrm{d}t} = -V_{\mathrm{Cell}} \cdot \left(\frac{[\mathrm{E}]\cdot \mathrm{k\_catC}\cdot [\mathrm{Met}]}{[\mathrm{Met}] + \mathrm{Km}_{(\mathrm{reaction\_10})}}\right)$$
$$+V_{\mathrm{Cell}} \cdot \left(\mathrm{k1}_{(\mathrm{reaction\_11})} \cdot [\mathrm{MetE}]\right)$$
$$-V_{\mathrm{Cell}} \cdot \left(\frac{[\mathrm{GlyE}]\cdot \mathrm{k\_cat}\cdot [\mathrm{Met}]}{[\mathrm{Met}] + \mathrm{Km}_{(\mathrm{reaction\_13})}}\right)$$
$$+V_{\mathrm{Cell}} \cdot \left(\mathrm{k1}_{(\mathrm{reaction\_16})} \cdot [\mathrm{EGM}]\right)$$
$$-V_{\mathrm{Cell}} \cdot (\mathrm{kInib} \cdot [\mathrm{SerE}] \cdot [\mathrm{Met}])$$
$$+V_{\mathrm{Cell}} \cdot (\mathrm{kInib} \cdot [\mathrm{ESM}])$$
$$-V_{\mathrm{Cell}} \cdot (\text{"kegm}+\text{"} \cdot [\mathrm{RnaGlyE}] \cdot [\mathrm{Met}])$$
$$+V_{\mathrm{Cell}} \cdot (\text{"kegm}-\text{"} \cdot [\mathrm{EGM}] \cdot [\mathrm{RNA}])$$
$$+V_{\mathrm{Cell}} \cdot \left(\mathrm{k1}_{(\mathrm{EST\_RNAinteraction2})} \cdot [\mathrm{ESTR}]\right)$$
$$-V_{\mathrm{Cell}} \cdot (\mathrm{kfull} \cdot [\mathrm{fRnaGlyE}] \cdot [\mathrm{Met}])$$
$$+V_{\mathrm{Cell}} \cdot (\mathrm{kfull} \cdot [\mathrm{EGM}] \cdot [\mathrm{fRNA}])$$

$$\frac{\mathrm{d}([\mathrm{Gly}]\cdot V_{\mathrm{Cell}})}{\mathrm{d}t} = -V_{\mathrm{Cell}}\cdot\left(\frac{[\mathrm{E}]\cdot \mathrm{k\_cat}\cdot[\mathrm{Gly}]}{[\mathrm{Gly}]+\mathrm{Km}_{(\mathrm{reaction\_9})}}\right)$$
$$+V_{\mathrm{Cell}}\cdot\left(\mathrm{k1}_{(\mathrm{reaction\_12})}\cdot[\mathrm{GlyE}]\right)$$
$$-V_{\mathrm{Cell}}\cdot\left(\frac{[\mathrm{MetE}]\cdot \mathrm{k\_cat}\cdot[\mathrm{Gly}]}{[\mathrm{Gly}]+\mathrm{Km}_{(\mathrm{reaction\_14})}}\right)$$
$$+V_{\mathrm{Cell}}\cdot\left(\mathrm{k1}_{(\mathrm{reaction\_15})}\cdot[\mathrm{EGM}]\right)$$
$$-V_{\mathrm{Cell}}\cdot(\mathrm{kInib}\cdot[\mathrm{EThf}]\cdot[\mathrm{Gly}])$$
$$+V_{\mathrm{Cell}}\cdot(\mathrm{kInib}\cdot[\mathrm{EGT}])$$
$$-V_{\mathrm{Cell}}\cdot(\mathrm{kg}\cdot[\mathrm{RNA\_E}]\cdot[\mathrm{Gly}])$$
$$+V_{\mathrm{Cell}}\cdot(\mathrm{kg}\cdot[\mathrm{RnaGlyE}])$$
$$-V_{\mathrm{Cell}}\cdot(\mathrm{K\_diff\_in}\cdot[\mathrm{Gly}])$$
$$+V_{\mathrm{Cell}}\cdot(\mathrm{k\_diff\_out}\cdot[\mathrm{Glym}])$$
$$+V_{\mathrm{Cell}}\cdot(\mathrm{alphaG})$$
$$-V_{\mathrm{Cell}}\cdot(\mathrm{betaG}\cdot[\mathrm{Gly}])$$
$$-V_{\mathrm{Cell}}\cdot(\mathrm{kfull}\cdot[\mathrm{fRNA\_E}]\cdot[\mathrm{Gly}])$$
$$+V_{\mathrm{Cell}}\cdot(\mathrm{kfull}\cdot[\mathrm{fRnaGlyE}])$$

$$\frac{\mathrm{d}([\mathrm{THF}]\cdot V_{\mathrm{Cell}})}{\mathrm{d}t} = -V_{\mathrm{Cell}}\cdot\left(\frac{[\mathrm{E}]\cdot \mathrm{k\_cat}_{(\mathrm{reaction})}\cdot[\mathrm{THF}]}{[\mathrm{THF}]+\mathrm{Km}_{(\mathrm{reaction})}}\right)$$
$$-V_{\mathrm{Cell}}\cdot\left(\frac{[\mathrm{SerE}]\cdot \mathrm{k\_cat}\cdot[\mathrm{THF}]}{[\mathrm{THF}]+\mathrm{Km}_{(\mathrm{reaction\_1})}}\right)$$
$$-V_{\mathrm{Cell}}\cdot(\mathrm{kInib}\cdot[\mathrm{EGly}]\cdot[\mathrm{THF}])$$
$$+V_{\mathrm{Cell}}\cdot(\mathrm{kInib}\cdot[\mathrm{EGT}])$$
$$-V_{\mathrm{Cell}}\cdot(\text{"kest}+\text{"}\cdot[\mathrm{RnaSerE}]\cdot[\mathrm{THF}])$$
$$+V_{\mathrm{Cell}}\cdot(\text{"kest}-\text{"}\cdot[\mathrm{EST}]\cdot[\mathrm{RNA}])$$
$$+V_{\mathrm{Cell}}\cdot\left(\mathrm{k1}_{(\mathrm{reaction\_4})}\cdot[\mathrm{EThf}]\right)$$
$$+V_{\mathrm{Cell}}\cdot\left(\mathrm{k1}_{(\mathrm{reaction\_6})}\cdot[\mathrm{EST}]\right)$$
$$-V_{\mathrm{Cell}}\cdot(\mathrm{kfull}\cdot[\mathrm{fRnaSerE}]\cdot[\mathrm{THF}])$$
$$+V_{\mathrm{Cell}}\cdot(\mathrm{kfull}\cdot[\mathrm{EST}]\cdot[\mathrm{fRNA}])$$

$$\begin{aligned}
\frac{\mathrm{d}([\mathrm{RNA}] \cdot V_{\mathrm{Cell}})}{\mathrm{d}t} = & -V_{\mathrm{Cell}} \cdot (\mathrm{kg} \cdot [\mathrm{RNA}] \cdot [\mathrm{E}]) \\
& + V_{\mathrm{Cell}} \cdot (\mathrm{kg} \cdot [\mathrm{RNA_E}]) \\
& - V_{\mathrm{Cell}} \cdot (\mathrm{kg} \cdot [\mathrm{RNA}] \cdot [\mathrm{SerE}]) \\
& + V_{\mathrm{Cell}} \cdot (\mathrm{kg} \cdot [\mathrm{RnaSerE}]) \\
& - V_{\mathrm{Cell}} \cdot (\mathrm{kg} \cdot [\mathrm{RNA}] \cdot [\mathrm{GlyE}]) \\
& + V_{\mathrm{Cell}} \cdot (\mathrm{kg} \cdot [\mathrm{RnaGlyE}]) \\
& + V_{\mathrm{Cell}} \cdot (\text{"kest + "} \cdot [\mathrm{RnaSerE}] \cdot [\mathrm{THF}]) \\
& - V_{\mathrm{Cell}} \cdot (\text{"kest − "} \cdot [\mathrm{EST}] \cdot [\mathrm{RNA}]) \\
& + V_{\mathrm{Cell}} \cdot (\text{"kegm + "} \cdot [\mathrm{RnaGlyE}] \cdot [\mathrm{Met}]) \\
& - V_{\mathrm{Cell}} \cdot (\text{"kegm − "} \cdot [\mathrm{EGM}] \cdot [\mathrm{RNA}]) \\
& - V_{\mathrm{Cell}} \cdot \left(\mathrm{k1}_{(\mathrm{RNA_degradation})} \cdot [\mathrm{RNA}]\right) \\
& - V_{\mathrm{Cell}} \cdot \left(\mathrm{k1}_{(\mathrm{EST_RNAinteraction})} \cdot [\mathrm{EST}] \cdot [\mathrm{RNA}]\right)
\end{aligned}$$

$$\begin{aligned}
\frac{\mathrm{d}([\mathrm{SerE}] \cdot V_{\mathrm{Cell}})}{\mathrm{d}t} = & + V_{\mathrm{Cell}} \cdot \left(\frac{[\mathrm{E}] \cdot \mathrm{k_cat} \cdot [\mathrm{Ser}]}{[\mathrm{Ser}] + \mathrm{Km}_{(\mathrm{reaction_0})}}\right) \\
& - V_{\mathrm{Cell}} \cdot (\mathrm{kInib} \cdot [\mathrm{SerE}] \cdot [\mathrm{Met}]) \\
& - V_{\mathrm{Cell}} \cdot \left(\frac{[\mathrm{SerE}] \cdot \mathrm{k_cat} \cdot [\mathrm{THF}]}{[\mathrm{THF}] + \mathrm{Km}_{(\mathrm{reaction_1})}}\right) \\
& + V_{\mathrm{Cell}} \cdot (\mathrm{kInib} \cdot [\mathrm{ESM}]) \\
& - V_{\mathrm{Cell}} \cdot (\mathrm{kg} \cdot [\mathrm{RNA}] \cdot [\mathrm{SerE}]) \\
& + V_{\mathrm{Cell}} \cdot (\mathrm{kg} \cdot [\mathrm{RnaSerE}]) \\
& - V_{\mathrm{Cell}} \cdot \left(\mathrm{k1}_{(\mathrm{reaction_3})} \cdot [\mathrm{SerE}]\right) \\
& - V_{\mathrm{Cell}} \cdot (\mathrm{kfull} \cdot [\mathrm{fRNA}] \cdot [\mathrm{SerE}]) \\
& + V_{\mathrm{Cell}} \cdot (\mathrm{kfull} \cdot [\mathrm{fRnaSerE}]) \\
& + V_{\mathrm{Cell}} \cdot \left(\mathrm{k1}_{(\mathrm{reaction_6})} \cdot [\mathrm{EST}]\right)
\end{aligned}$$

$$\frac{\mathrm{d}([\mathrm{GlyE}]\cdot V_{\mathrm{Cell}})}{\mathrm{d}t} = +V_{\mathrm{Cell}}\cdot\left(\frac{[\mathrm{E}]\cdot \mathrm{k\_cat}\cdot[\mathrm{Gly}]}{[\mathrm{Gly}]+\mathrm{Km}_{(\mathrm{reaction\_9})}}\right)$$
$$-V_{\mathrm{Cell}}\cdot\left(\mathrm{k1}_{(\mathrm{reaction\_12})}\cdot[\mathrm{GlyE}]\right)$$
$$-V_{\mathrm{Cell}}\cdot\left(\frac{[\mathrm{GlyE}]\cdot \mathrm{k\_cat}\cdot[\mathrm{Met}]}{[\mathrm{Met}]+\mathrm{Km}_{(\mathrm{reaction\_13})}}\right)$$
$$+V_{\mathrm{Cell}}\cdot\left(\mathrm{k1}_{(\mathrm{reaction\_16})}\cdot[\mathrm{EGM}]\right)$$
$$-V_{\mathrm{Cell}}\cdot(\mathrm{kg}\cdot[\mathrm{RNA}]\cdot[\mathrm{GlyE}])$$
$$+V_{\mathrm{Cell}}\cdot(\mathrm{kg}\cdot[\mathrm{RnaGlyE}])$$
$$-V_{\mathrm{Cell}}\cdot(\mathrm{kfull}\cdot[\mathrm{fRNA}]\cdot[\mathrm{GlyE}])$$
$$+V_{\mathrm{Cell}}\cdot(\mathrm{kfull}\cdot[\mathrm{fRnaGlyE}])$$

$$\frac{\mathrm{d}([\mathrm{EThf}]\cdot V_{\mathrm{Cell}})}{\mathrm{d}t} = +V_{\mathrm{Cell}}\cdot\left(\frac{[\mathrm{E}]\cdot \mathrm{k\_cat}_{(\mathrm{reaction})}\cdot[\mathrm{THF}]}{[\mathrm{THF}]+\mathrm{Km}_{(\mathrm{reaction})}}\right)$$
$$-V_{\mathrm{Cell}}\cdot(\mathrm{kInib}\cdot[\mathrm{EThf}]\cdot[\mathrm{Gly}])$$
$$+V_{\mathrm{Cell}}\cdot(\mathrm{kInib}\cdot[\mathrm{EGT}])$$
$$-V_{\mathrm{Cell}}\cdot\left(\frac{[\mathrm{EThf}]\cdot \mathrm{k\_cat}\cdot[\mathrm{Ser}]}{[\mathrm{Ser}]+\mathrm{Km}_{(\mathrm{reaction\_2})}}\right)$$
$$-V_{\mathrm{Cell}}\cdot\left(\mathrm{k1}_{(\mathrm{reaction\_4})}\cdot[\mathrm{EThf}]\right)$$
$$+V_{\mathrm{Cell}}\cdot\left(\mathrm{k1}_{(\mathrm{reaction\_5})}\cdot[\mathrm{EST}]\right)$$

$$\frac{\mathrm{d}([\mathrm{RNA\_E}]\cdot V_{\mathrm{Cell}})}{\mathrm{d}t} = +V_{\mathrm{Cell}}\cdot(\mathrm{kg}\cdot[\mathrm{RNA}]\cdot[\mathrm{E}])$$
$$-V_{\mathrm{Cell}}\cdot(\mathrm{kg}\cdot[\mathrm{RNA\_E}])$$
$$-V_{\mathrm{Cell}}\cdot(\mathrm{kg}\cdot[\mathrm{RNA\_E}]\cdot[\mathrm{Gly}])$$
$$+V_{\mathrm{Cell}}\cdot(\mathrm{kg}\cdot[\mathrm{RnaGlyE}])$$
$$-V_{\mathrm{Cell}}\cdot(\mathrm{kg}\cdot[\mathrm{RNA\_E}]\cdot[\mathrm{Ser}])$$
$$+V_{\mathrm{Cell}}\cdot\left(\mathrm{k1}_{(\mathrm{Rna\_10})}\cdot[\mathrm{RnaSerE}]\right)$$

$$\frac{\mathrm{d}([\mathrm{RnaSerE}]\cdot V_{\mathrm{Cell}})}{\mathrm{d}t} = +V_{\mathrm{Cell}}\cdot(\mathrm{kg}\cdot[\mathrm{RNA}]\cdot[\mathrm{SerE}])$$
$$-V_{\mathrm{Cell}}\cdot(\mathrm{kg}\cdot[\mathrm{RnaSerE}])$$
$$+V_{\mathrm{Cell}}\cdot(\mathrm{kg}\cdot[\mathrm{RNA\_E}]\cdot[\mathrm{Ser}])$$
$$-V_{\mathrm{Cell}}\cdot\left(\mathrm{k1}_{(\mathrm{Rna\_10})}\cdot[\mathrm{RnaSerE}]\right)$$
$$-V_{\mathrm{Cell}}\cdot(\text{"kest + "}\cdot[\mathrm{RnaSerE}]\cdot[\mathrm{THF}])$$
$$+V_{\mathrm{Cell}}\cdot(\text{"kest − "}\cdot[\mathrm{EST}]\cdot[\mathrm{RNA}])$$

$$\frac{\mathrm{d}([\mathrm{RnaGlyE}]\cdot V_{\mathrm{Cell}})}{\mathrm{d}t} = +V_{\mathrm{Cell}}\cdot(\mathrm{kg}\cdot[\mathrm{RNA}]\cdot[\mathrm{GlyE}])$$
$$-V_{\mathrm{Cell}}\cdot(\mathrm{kg}\cdot[\mathrm{RnaGlyE}])$$
$$+V_{\mathrm{Cell}}\cdot(\mathrm{kg}\cdot[\mathrm{RNA\_E}]\cdot[\mathrm{Gly}])$$
$$-V_{\mathrm{Cell}}\cdot(\mathrm{kg}\cdot[\mathrm{RnaGlyE}])$$
$$-V_{\mathrm{Cell}}\cdot(\text{"kegm + "}\cdot[\mathrm{RnaGlyE}]\cdot[\mathrm{Met}])$$
$$+V_{\mathrm{Cell}}\cdot(\text{"kegm − "}\cdot[\mathrm{EGM}]\cdot[\mathrm{RNA}])$$
$$+V_{\mathrm{Cell}}\cdot\left(\mathrm{k1}_{(\mathrm{EST\_RNAinteraction2})}\cdot[\mathrm{ESTR}]\right)$$

$$\frac{\mathrm{d}([\mathrm{EST}]\cdot V_{\mathrm{Cell}})}{\mathrm{d}t} = +V_{\mathrm{Cell}}\cdot\left(\frac{[\mathrm{SerE}]\cdot\mathrm{k\_cat}\cdot[\mathrm{THF}]}{[\mathrm{THF}]+\mathrm{Km}_{(\mathrm{reaction\_1})}}\right)$$
$$+V_{\mathrm{Cell}}\cdot\left(\frac{[\mathrm{EThf}]\cdot\mathrm{k\_cat}\cdot[\mathrm{Ser}]}{[\mathrm{Ser}]+\mathrm{Km}_{(\mathrm{reaction\_2})}}\right)$$
$$+V_{\mathrm{Cell}}\cdot(\text{"kest + "}\cdot[\mathrm{RnaSerE}]\cdot[\mathrm{THF}])$$
$$-V_{\mathrm{Cell}}\cdot(\text{"kest − "}\cdot[\mathrm{EST}]\cdot[\mathrm{RNA}])$$
$$-V_{\mathrm{Cell}}\cdot\left(\mathrm{k1}_{(\mathrm{EST\_RNAinteraction})}\cdot[\mathrm{EST}]\cdot[\mathrm{RNA}]\right)$$
$$-V_{\mathrm{Cell}}\cdot\left(\mathrm{k1}_{(\mathrm{reaction\_5})}\cdot[\mathrm{EST}]\right)$$
$$-V_{\mathrm{Cell}}\cdot\left(\mathrm{k1}_{(\mathrm{reaction\_6})}\cdot[\mathrm{EST}]\right)$$
$$+V_{\mathrm{Cell}}\cdot(\mathrm{kfull}\cdot[\mathrm{fRnaSerE}]\cdot[\mathrm{THF}])$$
$$-V_{\mathrm{Cell}}\cdot(\mathrm{kfull}\cdot[\mathrm{EST}]\cdot[\mathrm{fRNA}])$$
$$-V_{\mathrm{Cell}}\cdot(\mathrm{k\_catC}\cdot[\mathrm{EST}])$$
$$+V_{\mathrm{Cell}}\cdot(\text{"k\_catC − "}\cdot[\mathrm{EGM}])$$

$$\frac{\mathrm{d}([\mathrm{EGM}]\cdot V_{\mathrm{Cell}})}{\mathrm{d}t} = +V_{\mathrm{Cell}}\cdot\left(\frac{[\mathrm{GlyE}]\cdot\mathrm{k\_cat}\cdot[\mathrm{Met}]}{[\mathrm{Met}]+\mathrm{Km}_{(\mathrm{reaction\_13})}}\right)$$
$$+V_{\mathrm{Cell}}\cdot\left(\frac{[\mathrm{MetE}]\cdot\mathrm{k\_cat}\cdot[\mathrm{Gly}]}{[\mathrm{Gly}]+\mathrm{Km}_{(\mathrm{reaction\_14})}}\right)$$
$$-V_{\mathrm{Cell}}\cdot\left(\mathrm{k1}_{(\mathrm{reaction\_15})}\cdot[\mathrm{EGM}]\right)$$
$$-V_{\mathrm{Cell}}\cdot\left(\mathrm{k1}_{(\mathrm{reaction\_16})}\cdot[\mathrm{EGM}]\right)$$
$$+V_{\mathrm{Cell}}\cdot(\text{"kegm + "}\cdot[\mathrm{RnaGlyE}]\cdot[\mathrm{Met}])$$
$$-V_{\mathrm{Cell}}\cdot(\text{"kegm − "}\cdot[\mathrm{EGM}]\cdot[\mathrm{RNA}])$$
$$+V_{\mathrm{Cell}}\cdot(\mathrm{kfull}\cdot[\mathrm{fRnaGlyE}]\cdot[\mathrm{Met}])$$
$$-V_{\mathrm{Cell}}\cdot(\mathrm{kfull}\cdot[\mathrm{EGM}]\cdot[\mathrm{fRNA}])$$
$$+V_{\mathrm{Cell}}\cdot(\mathrm{k\_catC}\cdot[\mathrm{EST}])$$
$$-V_{\mathrm{Cell}}\cdot(\text{"k\_catC − "}\cdot[\mathrm{EGM}])$$

$$\frac{\mathrm{d}([\mathrm{MetE}] \cdot V_{\mathrm{Cell}})}{\mathrm{d}t} = + V_{\mathrm{Cell}} \cdot \left(\frac{[\mathrm{E}] \cdot \mathrm{k\_catC} \cdot [\mathrm{Met}]}{[\mathrm{Met}] + \mathrm{Km}_{(\mathrm{reaction\_10})}}\right)$$
$$- V_{\mathrm{Cell}} \cdot \left(\mathrm{k1}_{(\mathrm{reaction\_11})} \cdot [\mathrm{MetE}]\right)$$
$$- V_{\mathrm{Cell}} \cdot \left(\frac{[\mathrm{MetE}] \cdot \mathrm{k\_cat} \cdot [\mathrm{Gly}]}{[\mathrm{Gly}] + \mathrm{Km}_{(\mathrm{reaction\_14})}}\right)$$
$$+ V_{\mathrm{Cell}} \cdot \left(\mathrm{k1}_{(\mathrm{reaction\_15})} \cdot [\mathrm{EGM}]\right)$$
$$- V_{\mathrm{Cell}} \cdot (\mathrm{kInib} \cdot [\mathrm{MetE}] \cdot [\mathrm{Ser}])$$
$$+ V_{\mathrm{Cell}} \cdot (\mathrm{kInib} \cdot [\mathrm{ESM}])$$

$$\frac{\mathrm{d}([\mathrm{ESM}] \cdot V_{\mathrm{Cell}})}{\mathrm{d}t} = + V_{\mathrm{Cell}} \cdot (\mathrm{kInib} \cdot [\mathrm{MetE}] \cdot [\mathrm{Ser}])$$
$$+ V_{\mathrm{Cell}} \cdot (\mathrm{kInib} \cdot [\mathrm{SerE}] \cdot [\mathrm{Met}])$$
$$- V_{\mathrm{Cell}} \cdot (\mathrm{kInib} \cdot [\mathrm{ESM}])$$
$$- V_{\mathrm{Cell}} \cdot (\mathrm{kInib} \cdot [\mathrm{ESM}])$$

$$\frac{\mathrm{d}([\mathrm{EGT}] \cdot V_{\mathrm{Cell}})}{\mathrm{d}t} = + V_{\mathrm{Cell}} \cdot (\mathrm{kInib} \cdot [\mathrm{EThf}] \cdot [\mathrm{Gly}])$$
$$+ V_{\mathrm{Cell}} \cdot (\mathrm{kInib} \cdot [\mathrm{EGly}] \cdot [\mathrm{THF}])$$
$$- V_{\mathrm{Cell}} \cdot (\mathrm{kInib} \cdot [\mathrm{EGT}])$$
$$- V_{\mathrm{Cell}} \cdot (\mathrm{kInib} \cdot [\mathrm{EGT}])$$

$$\frac{\mathrm{d}([\mathrm{EGly}] \cdot V_{\mathrm{Cell}})}{\mathrm{d}t} = - V_{\mathrm{Cell}} \cdot (\mathrm{kInib} \cdot [\mathrm{EGly}] \cdot [\mathrm{THF}])$$
$$+ V_{\mathrm{Cell}} \cdot (\mathrm{kInib} \cdot [\mathrm{EGT}])$$

$$\frac{\mathrm{d}([\mathrm{Em}] \cdot V_{\mathrm{Cell}})}{\mathrm{d}t} = + V_{\mathrm{Cell}} \cdot (\mathrm{K\_diff\_in} \cdot [\mathrm{EmC}])$$
$$- V_{\mathrm{Cell}} \cdot (\mathrm{mu} \cdot [\mathrm{Em}])$$
$$- V_{\mathrm{Cell}} \cdot \left(\frac{[\mathrm{Em}] \cdot \mathrm{k\_cat} \cdot [\mathrm{Serm}]}{[\mathrm{Serm}] + \mathrm{Km}_{(\mathrm{mitocondria\_1})}}\right)$$
$$+ V_{\mathrm{Cell}} \cdot \left(\mathrm{k1}_{(\mathrm{mitocondria\_2})} \cdot [\mathrm{SerEm}]\right)$$
$$- V_{\mathrm{Cell}} \cdot \left(\frac{[\mathrm{Em}] \cdot \mathrm{k\_cat} \cdot [\mathrm{THFm}]}{[\mathrm{THFm}] + \mathrm{Km}_{(\mathrm{mitocondria\_4})}}\right)$$
$$+ V_{\mathrm{Cell}} \cdot \left(\mathrm{k1}_{(\mathrm{mitocondria\_5})} \cdot [\mathrm{EThfm}]\right)$$
$$+ V_{\mathrm{Cell}} \cdot \left(\mathrm{k1}_{(\mathrm{mitocondria\_6})} \cdot [\mathrm{EGMm}]\right)$$
$$- V_{\mathrm{Cell}} \cdot \left(\mathrm{k1}_{(\mathrm{mitocondria\_7})} \cdot [\mathrm{Glym}] \cdot [\mathrm{Metm}] \cdot [\mathrm{Em}]\right)$$

$$\frac{\mathrm{d}([\mathrm{EmC}] \cdot V_{\mathrm{Cell}})}{\mathrm{d}t} = - V_{\mathrm{Cell}} \cdot (\mathrm{K\_diff\_in} \cdot [\mathrm{EmC}])$$
$$+ V_{\mathrm{Cell}} \cdot \left(\frac{\mathrm{kTransl} \cdot [\mathrm{RNA}]}{[\mathrm{RNA}] + K_{(\mathrm{RNA\_translation})}}\right)$$

$$\frac{\mathrm{d}([\mathrm{Serm}]\cdot V_{\mathrm{Cell}})}{\mathrm{d}t} = +V_{\mathrm{Cell}} \cdot (\mathrm{K\_diff\_in} \cdot [\mathrm{Ser}]) - V_{\mathrm{Cell}} \cdot (\mathrm{k\_diff\_out} \cdot [\mathrm{Serm}]) - V_{\mathrm{Cell}} \cdot \left(\frac{[\mathrm{Em}]\cdot \mathrm{k\_cat}\cdot[\mathrm{Serm}]}{[\mathrm{Serm}]+\mathrm{Km}_{(\mathrm{mitocondria\_1})}}\right) + V_{\mathrm{Cell}} \cdot \left(\mathrm{k1}_{(\mathrm{mitocondria\_2})} \cdot [\mathrm{SerEm}]\right)$$

$$\frac{\mathrm{d}([\mathrm{Glym}]\cdot V_{\mathrm{Cell}})}{\mathrm{d}t} = +V_{\mathrm{Cell}} \cdot (\mathrm{K\_diff\_in} \cdot [\mathrm{Gly}]) - V_{\mathrm{Cell}} \cdot (\mathrm{k\_diff\_out} \cdot [\mathrm{Glym}]) + V_{\mathrm{Cell}} \cdot \left(\mathrm{k1}_{(\mathrm{mitocondria\_6})} \cdot [\mathrm{EGMm}]\right) - V_{\mathrm{Cell}} \cdot \left(\mathrm{k1}_{(\mathrm{mitocondria\_7})} \cdot [\mathrm{Glym}] \cdot [\mathrm{Metm}] \cdot [\mathrm{Em}]\right) - V_{\mathrm{Cell}} \cdot (\mathrm{k\_cons} \cdot [\mathrm{Glym}])$$

$$\frac{\mathrm{d}([\mathrm{SerEm}]\cdot V_{\mathrm{Cell}})}{\mathrm{d}t} = +V_{\mathrm{Cell}} \cdot \left(\frac{[\mathrm{Em}]\cdot \mathrm{k\_cat}\cdot[\mathrm{Serm}]}{[\mathrm{Serm}]+\mathrm{Km}_{(\mathrm{mitocondria\_1})}}\right) - V_{\mathrm{Cell}} \cdot \left(\mathrm{k1}_{(\mathrm{mitocondria\_2})} \cdot [\mathrm{SerEm}]\right) - V_{\mathrm{Cell}} \cdot \left(\frac{[\mathrm{SerEm}]\cdot \mathrm{k\_cat}\cdot[\mathrm{THFm}]}{[\mathrm{THFm}]+\mathrm{Km}_{(\mathrm{mitocondria\_3})}}\right)$$

$$\frac{\mathrm{d}([\mathrm{EGMm}]\cdot V_{\mathrm{Cell}})}{\mathrm{d}t} = -V_{\mathrm{Cell}} \cdot \left(\mathrm{k1}_{(\mathrm{mitocondria\_6})} \cdot [\mathrm{EGMm}]\right) + V_{\mathrm{Cell}} \cdot \left(\mathrm{k1}_{(\mathrm{mitocondria\_7})} \cdot [\mathrm{Glym}] \cdot [\mathrm{Metm}] \cdot [\mathrm{Em}]\right) + V_{\mathrm{Cell}} \cdot (\mathrm{k\_catM} \cdot [\mathrm{ESTm}])$$

$$\frac{\mathrm{d}([\mathrm{THFm}]\cdot V_{\mathrm{Cell}})}{\mathrm{d}t} = -V_{\mathrm{Cell}} \cdot \left(\frac{[\mathrm{SerEm}]\cdot \mathrm{k\_cat}\cdot[\mathrm{THFm}]}{[\mathrm{THFm}]+\mathrm{Km}_{(\mathrm{mitocondria\_3})}}\right) - V_{\mathrm{Cell}} \cdot \left(\frac{[\mathrm{Em}]\cdot \mathrm{k\_cat}\cdot[\mathrm{THFm}]}{[\mathrm{THFm}]+\mathrm{Km}_{(\mathrm{mitocondria\_4})}}\right) + V_{\mathrm{Cell}} \cdot \left(\mathrm{k1}_{(\mathrm{mitocondria\_5})} \cdot [\mathrm{EThfm}]\right)$$

$$\frac{\mathrm{d}([\mathrm{EThfm}]\cdot V_{\mathrm{Cell}})}{\mathrm{d}t} = +V_{\mathrm{Cell}} \cdot \left(\frac{[\mathrm{Em}]\cdot \mathrm{k\_cat}\cdot[\mathrm{THFm}]}{[\mathrm{THFm}]+\mathrm{Km}_{(\mathrm{mitocondria\_4})}}\right) - V_{\mathrm{Cell}} \cdot \left(\mathrm{k1}_{(\mathrm{mitocondria\_5})} \cdot [\mathrm{EThfm}]\right)$$

$$\frac{\mathrm{d}([\mathrm{Metm}]\cdot V_{\mathrm{Cell}})}{\mathrm{d}t} = +V_{\mathrm{Cell}} \cdot \left(\mathrm{k1}_{(\mathrm{mitocondria\_6})} \cdot [\mathrm{EGMm}]\right) - V_{\mathrm{Cell}} \cdot \left(\mathrm{k1}_{(\mathrm{mitocondria\_7})} \cdot [\mathrm{Glym}] \cdot [\mathrm{Metm}] \cdot [\mathrm{Em}]\right)$$

$$\frac{\mathrm{d}([\mathrm{ESTR}]\cdot V_{\mathrm{Cell}})}{\mathrm{d}t} = +V_{\mathrm{Cell}} \cdot \left(\mathrm{k1}_{(\mathrm{EST\_RNAinteraction})} \cdot [\mathrm{EST}] \cdot [\mathrm{RNA}]\right) - V_{\mathrm{Cell}} \cdot \left(\mathrm{k1}_{(\mathrm{EST\_RNAinteraction2})} \cdot [\mathrm{ESTR}]\right)$$

$$\frac{\mathrm{d}([\mathrm{ESTm}]\cdot V_{\mathrm{Cell}})}{\mathrm{d}t} = +V_{\mathrm{Cell}} \cdot \left(\frac{[\mathrm{SerEm}]\cdot \mathrm{k\_cat}\cdot[\mathrm{THFm}]}{[\mathrm{THFm}]+\mathrm{Km}_{(\mathrm{mitocondria\_3})}}\right) - V_{\mathrm{Cell}} \cdot (\mathrm{k\_catM} \cdot [\mathrm{ESTm}])$$

$$
\begin{aligned}
\frac{\mathrm{d}([\text{fRNA}] \cdot V_{\text{Cell}})}{\mathrm{d}t} = & -V_{\text{Cell}} \cdot (\text{kfull} \cdot [\text{fRNA}] \cdot [\text{E}]) \\
& +V_{\text{Cell}} \cdot (\text{kfull} \cdot [\text{fRNA\_E}]) \\
& -V_{\text{Cell}} \cdot (\text{kfull} \cdot [\text{fRNA}] \cdot [\text{SerE}]) \\
& +V_{\text{Cell}} \cdot (\text{kfull} \cdot [\text{fRnaSerE}]) \\
& -V_{\text{Cell}} \cdot (\text{kfull} \cdot [\text{fRNA}] \cdot [\text{GlyE}]) \\
& +V_{\text{Cell}} \cdot (\text{kfull} \cdot [\text{fRnaGlyE}]) \\
& +V_{\text{Cell}} \cdot (\text{kfull} \cdot [\text{fRnaSerE}] \cdot [\text{THF}]) \\
& -V_{\text{Cell}} \cdot (\text{kfull} \cdot [\text{EST}] \cdot [\text{fRNA}]) \\
& +V_{\text{Cell}} \cdot (\text{kfull} \cdot [\text{fRnaGlyE}] \cdot [\text{Met}]) \\
& -V_{\text{Cell}} \cdot (\text{kfull} \cdot [\text{EGM}] \cdot [\text{fRNA}])
\end{aligned}
$$

$$
\begin{aligned}
\frac{\mathrm{d}([\text{fRNA\_E}] \cdot V_{\text{Cell}})}{\mathrm{d}t} = & +V_{\text{Cell}} \cdot (\text{kfull} \cdot [\text{fRNA}] \cdot [\text{E}]) \\
& -V_{\text{Cell}} \cdot (\text{kfull} \cdot [\text{fRNA\_E}]) \\
& -V_{\text{Cell}} \cdot (\text{kfull} \cdot [\text{fRNA\_E}] \cdot [\text{Gly}]) \\
& +V_{\text{Cell}} \cdot (\text{kfull} \cdot [\text{fRnaGlyE}]) \\
& -V_{\text{Cell}} \cdot (\text{kfull} \cdot [\text{fRNA\_E}] \cdot [\text{Ser}]) \\
& +V_{\text{Cell}} \cdot (\text{kfull} \cdot [\text{fRnaSerE}])
\end{aligned}
$$

$$
\begin{aligned}
\frac{\mathrm{d}([\text{fRnaSerE}] \cdot V_{\text{Cell}})}{\mathrm{d}t} = & +V_{\text{Cell}} \cdot (\text{kfull} \cdot [\text{fRNA}] \cdot [\text{SerE}]) \\
& -V_{\text{Cell}} \cdot (\text{kfull} \cdot [\text{fRnaSerE}]) \\
& +V_{\text{Cell}} \cdot (\text{kfull} \cdot [\text{fRNA\_E}] \cdot [\text{Ser}]) \\
& -V_{\text{Cell}} \cdot (\text{kfull} \cdot [\text{fRnaSerE}]) \\
& -V_{\text{Cell}} \cdot (\text{kfull} \cdot [\text{fRnaSerE}] \cdot [\text{THF}]) \\
& +V_{\text{Cell}} \cdot (\text{kfull} \cdot [\text{EST}] \cdot [\text{fRNA}])
\end{aligned}
$$

$$
\begin{aligned}
\frac{\mathrm{d}([\text{fRnaGlyE}] \cdot V_{\text{Cell}})}{\mathrm{d}t} = & +V_{\text{Cell}} \cdot (\text{kfull} \cdot [\text{fRNA}] \cdot [\text{GlyE}]) \\
& -V_{\text{Cell}} \cdot (\text{kfull} \cdot [\text{fRnaGlyE}]) \\
& +V_{\text{Cell}} \cdot (\text{kfull} \cdot [\text{fRNA\_E}] \cdot [\text{Gly}]) \\
& -V_{\text{Cell}} \cdot (\text{kfull} \cdot [\text{fRnaGlyE}]) \\
& -V_{\text{Cell}} \cdot (\text{kfull} \cdot [\text{fRnaGlyE}] \cdot [\text{Met}]) \\
& +V_{\text{Cell}} \cdot (\text{kfull} \cdot [\text{EGM}] \cdot [\text{fRNA}])
\end{aligned}
$$

#### V. REACTION TABLE

Here we present the table of the reactions involved with their relative rate law, that are defined in the differential equations presented above. Shown here is also the table of the physical constants with their dimension and their values.

| Reaction Name | Definition | Rate Law |
|---|---|---|
| OCM reaction 0 | E + Ser → SerE; Ser E | Saturation Reaction |
| OCM reaction 1 | E + THF → EThf; THF E | Saturation Reaction |
| OCM reaction 2 | SerE + THF → EST; THF SerE | Saturation Reaction |
| OCM reaction 3 | EThf + Ser → EST; Ser EThf | Saturation Reaction |
| OCM reaction 4 | SerE → E + Ser | Mass Action |
| OCM reaction 5 | EThf → E + THF | Mass Action |
| OCM reaction 6 | EST → Ser + EThf | Mass Action |
| OCM reaction 7 | EST → SerE + THF | Mass Action |
| OCM reaction 8 | EST → EGM | Mass Action |
| OCM reaction 9 | EGM → EST | Mass Action |
| OCM reaction 10 | E + Gly → GlyE; Gly E | Saturation Reaction |
| OCM reaction 11 | E + Met → MetE; Met E | Saturation Reaction |
| OCM reaction 13 | MetE → E + Met | Mass Action |
| OCM reaction 14 | GlyE → E + Gly | Mass Action |
| OCM reaction 15 | GlyE + Met → EGM; Met GlyE | Mass Action |
| OCM reaction 16 | MetE + Gly → EGM; Gly MetE | Mass Action |
| OCM reaction 17 | EGM → MetE + Gly | Mass Action |
| OCM reaction 18 | EGM → GlyE + Met | Mass Action |
| Inhibition reaction 0 | MetE + Ser → ESM; Ser MetE | Mass Action |
| Inhibition reaction 1 | SerE + Met → ESM | Mass Action |
| Inhibition reaction 2 | ESM → SerE + Met | Mass Action |
| Inhibition reaction 3 | ESM → MetE + Ser | Mass Action |
| Complex formation reaction 0 | EThf + Gly → EGT | Mass Action |
| Complex formation reaction 1 | EGly + THF → EGT | Mass Action |
| Complex formation reaction 2 | EGT → EThf + Gly | Mass Action |
| Complex formation reaction 3 | EGT → EGly + THF | Mass Action |

TABLE I: One carbon metabolism (OCM) reactions involved, free RNAs system

| Reaction Name | Definition | Constants |
|---|---|---|
| RNA reactions 0 | RNA + E → RNA_E | Mass Action |
| RNA reactions 1 | RNA_E → RNA + E | Mass Action |
| RNA reactions 2 | RNA + SerE → RnaSerE | Mass Action |
| RNA reactions 3 | RnaSerE → RNA + SerE | Mass Action |
| RNA reactions 4 | RNA + GlyE → RnaGlyE | Mass Action |
| RNA reactions 5 | RnaGlyE → RNA + GlyE | Mass Action |
| RNA reactions 6 | RNA_E + Gly → RnaGlyE | Mass Action |
| RNA reactions 7 | RnaGlyE → RNA_E + Gly | Mass Action |
| RNA reactions 8 | RNA_E + Ser → RnaSerE | Mass Action |
| RNA reactions 9 | RnaSerE → RNA_E + Ser | Mass Action |
| RNA reactions 10 | RnaSerE + THF → EST + RNA | Mass Action |
| RNA reactions 11 | EST + RNA → RnaSerE + THF | Mass Action |
| RNA reactions 12 | RnaGlyE + Met → EGM + RNA | Mass Action |
| RNA reactions 13 | EGM + RNA → RnaGlyE + Met | Mass Action |
| RNA degradation | → RNA | Mass Action |
| RNA expression | RNA → | Mass Action |
| EST_RNAinteraction | EST + RNA → ESTR | Mass Action |
| EST_RNAinteraction 2 | ESTR → RnaGlyE + Met | Mass Action |
| RNA translation | RNA → EmC + RNA; RNA | Translation |

TABLE II: UTR2 RNA reactions involved

| Reaction Name | Definition | Rate Law |
|---|---|---|
| SHMT2 transport mitochondrion | EmC → Em | Mass Action |
| Mit Glicine degradation | Glym → | Mass Action |
| Mit Serine degradation | Serm → | Mass Action |
| Glycine creation cytosol | → Gly | Mass Action |
| Serine creation cytosol | → Ser | Mass Action |
| Glycine degradation cytosol | Gly → | Mass Action |
| Serine degradation cytosol | Ser → | Mass Action |
| SHMT 2 degradation | Em → | Mass Action |
| Serine Transport to Mit | Ser → Serm | Mass Action |
| Glycine Transport to Mit | Gly → Glym | Mass Action |
| Serine Transport to cytosol | Serm → Ser | Mass Action |
| Glycine Transport to cytosol | Glym → Gly | Mass Action |
| Mitochondrion Reaction 1 | Serm + Em → SerEm; Serm Em | Saturation Reaction |
| Mitochondrion Reaction 2 | SerEm → Serm + Em | Mass Action |
| Mitochondrion Reaction 3 | SerEm + THFm → ESTm; THFm SerEm | Saturation Reaction |
| Mitochondrion Reaction 4 | Em + THFm → EThfm; THFm Em | Mass Action |
| Mitochondrion Reaction 5 | EThfm → Em + THFm | Saturation Reaction |
| Mitochondrion Reaction 6 | EGMm → Glym + Metm + Em | Mass Action |
| Mitochondrion Reaction 7 | Glym + Metm + Em → EGMm | Mass Action |

TABLE III: Mitochondrial, transport, production and degradation reactions

| Reaction Name | Definition | Rate Law |
|---|---|---|
| poolRNA reaction 0 | fRNA + E → fRNA_E | Mass Action |
| poolRNA reaction 1 | fRNA_E → fRNA + E | Mass Action |
| poolRNA reaction 3 | fRNA + SerE → fRnaSerE | Mass Action |
| poolRNA reaction 4 | fRnaSerE → fRNA + SerE | Mass Action |
| poolRNA reaction 5 | fRNA + GlyE → fRnaGlyE | Mass Action |
| poolRNA reaction 6 | fRnaGlyE → fRNA + GlyE | Mass Action |
| poolRNA reaction 7 | fRNA_E + Gly → fRnaGlyE | Mass Action |
| poolRNA reaction 8 | fRnaGlyE → fRNA_E + Gly | Mass Action |
| poolRNA reaction 9 | fRNA_E + Ser → fRnaSerE | Mass Action |
| poolRNA reaction 10 | fRnaSerE → fRNA_E + Ser | Mass Action |
| poolRNA reaction 11 | fRnaSerE + THF → EST + fRNA | Mass Action |
| poolRNA reaction 12 | EST + fRNA → fRnaSerE + THF | Mass Action |
| poolRNA reaction 13 | fRnaGlyE + Met → EGM + fRNA | Mass Action |
| poolRNA reaction 14 | EGM + fRNA → fRnaGlyE + Met | Mass Action |

TABLE IV: RNApool reactions involved

| Symbol | Molecular name | Compartment |
|---|---|---|
| E | SHMT1 | Cytosol |
| Ser | Serine | Cytosol |
| Met | methylentetrahydrofolate | Cytosol |
| Gly | Glycine | Cytosol |
| THF | Tetrahydrofolate | Cytosol |
| RNA | mRNA SHMT2 (UTR2) | Cytosol |
| SerE | Serine-SHMT1 complex | Cytosol |
| GlyE | Glycine-SHMT1 complex | Cytosol |
| EThf | SHMT1 tetrafolate complex | Cytosol |
| RNA_E | UTR2 - SHMT1 complex | Cytosol |
| RnaSerE | UTR2 - SHMT1 - Serine complex | Cytosol |
| RnaGlyE | UTR2 - SHMT1 - Glycine complex | Cytosol |
| EST | SHMT1 - Serine - THF complex | Cytosol |
| EGM | SHMT1 - Glycine - Met complex | Cytosol |
| MetE | SHMT1 - Met complex | Cytosol |
| ESM | SHMT1 - Serine- Met complex | Cytosol |
| EGT | SHMT1 - Glycine - THF complex | Cytosol |
| EGly | SHMT1 - glycine complex | Cytosol |
| Em | SHMT2 | Mitochondrion |
| EmC | SHMT2 | Cytosol |
| Serm | Serine | Mitochondrion |
| Glym | Glycine | Mitochondrion |
| SerEm | SHMT2 - Serine complex | Mitochondrion |
| EGMm | SHMT2 - Glycine - Met complex | Mitochondrion |
| THFm | Tetrahydrofolate | Mitochondrion |
| EThfm | SHMT2 - THF complex | Mitochondrion |
| Metm | methylentetrahydrofolate | Mitochondrion |
| ESTR | SHMT1 - Serine - THF - UTR2 complex | Cytosol |
| ESTm | SHMT2 - Serine - THF complex | Mitochondrion |
| fRNA | RNApool | Cytosol |
| fRNA_E | RNApool - SHMT1 complex | Cytosol |
| fRnaSerE | RNApool - SHMT1- Glycine complex | Cytosol |
| fRnaGlyE | RNApool - SHMT1 - serine complex | Cytosol |

TABLE V: Legend of the symbols used in the paper

| Constant Name | Value | Physical dimension |
|---|---|---|
| k1 | 0.1 | $time^{-1}$ |
| kg | 0.1 | $time^{-1}$ |
| kest+ | 0.0001 | $time^{-1}$ |
| kest- | 0.1 | $time^{-1}$ |
| kegm+ | 0.1 | $time^{-1}$ |
| kegm- | 0.1 | $time^{-1}$ |
| k_diff_out | 0.0001 | $time^{-1}$ |
| K_diff_in | 0.0001 | $time^{-1}$ |
| kTransl | 0.1 | $time^{-1}$ |
| mu | 10 | $time^{-1}$ |
| k_catC | 374 | $time^{-1}$ |
| k_catM | 813 | $time^{-1}$ |
| k_cat | 100 | $time^{-1}$ |
| k_catC- | 350 | $time^{-1}$ |
| kInib | 0.1 | $time^{-1}$ |
| k_full | 0.1 | $time^{-1}$ |
| k_cons | 0.1 | $time^{-1}$ |
| alphaG | 0.00001 | $time^{-1}$ |
| betaG | 0.001 | $time^{-1}$ |
| alphaS | 0.00001 | $time^{-1}$ |
| betaS | 0.001 | $time^{-1}$ |
| Km0 | 0.033 | concentration |
| Km1 | 4.8 | concentration |
| Km2 | 0.007 | concentration |
| Km10 | 0.084 | concentration |
| Vcell | 1 | volume |

TABLE VI: Table of physical constants used

[1] Daniel T. Gillespie. Exact stochastic simulation of coupled chemical reactions. *The Journal of Physical Chemistry*, 81(25):2340–2361, 1977.

[2] M. Bellucci, F. Agostini, M. Masin, and G. G. Tartaglia. Predicting protein associations with long noncoding RNAs. *Nat Methods*, 8(6):444–445, Jun 2011.

[3] B. Lang, A. Armaos, and G. G. Tartaglia. RNAct: Protein-RNA interaction predictions for model organisms with supporting experimental data. *Nucleic Acids Res*, 47(D1):D601–D606, 01 2019.
